# The Integrated Stress Response Suppresses PINK1-dependent Mitophagy by Preserving Mitochondrial Import Efficiency

**DOI:** 10.1101/2024.10.16.617214

**Authors:** Mingchong Yang, Zengshuo Mo, Kelly Walsh, Wen Liu, Xiaoyan Guo

**Affiliations:** Department of Genetics and Genome Sciences, University of Connecticut Health Center, Farmington, CT, 06032; Department of Molecular Biology and Biophysics, University of Connecticut Health Center, Farmington, CT, 06032

**Keywords:** Integrated stress response (ISR), mitophagy, mitochondrial protein import

## Abstract

Mitophagy is crucial for maintaining mitochondrial health, but how its levels adjust to different stress conditions remains unclear. In this study, we investigated the role of the DELE1-HRI axis of integrated stress response (ISR) in regulating mitophagy, a key mitochondrial stress pathway. Our findings show that the ISR suppresses mitophagy under non-depolarizing mitochondrial stress by positively regulating mitochondrial protein import, independent of ATF4 activation. Mitochondrial protein import is regulated by the rate of protein synthesis under both depolarizing and non-depolarizing stress.

Without ISR, increased protein synthesis overwhelms the mitochondrial import machinery, reducing its efficiency. Under depolarizing stress, mitochondrial import is heavily impaired even with active ISR, leading to significant PINK1 accumulation. In contrast, non-depolarizing stress allows more efficient protein import in the presence of ISR, resulting in lower mitophagy. Without ISR, mitochondrial protein import becomes severely compromised, causing PINK1 accumulation to reach the threshold necessary to trigger mitophagy. These findings reveal a novel link between ISR-regulated protein synthesis, mitochondrial import, and mitophagy, offering potential therapeutic targets for diseases associated with mitochondrial dysfunction.

## Introduction

Mitochondria, best known as the powerhouse of the cell, play crucial roles beyond energy production, including metabolic regulation, calcium homeostasis, and apoptosis [1]. To maintain mitochondrial function and cellular homeostasis, mitochondria engage in extensive crosstalk with other cellular components [2]. One key molecular pathway activated in response to many, if not all mitochondrial stress conditions, is the integrated stress response (ISR) [3, 4]. The ISR is a general adaptive mechanism that cells use to cope with a broad range of stress conditions, including viral infections, amino acid deprivation, heme depletion, and endoplasmic reticulum (ER) stress [5]. While ISR pathways are triggered by distinct stimuli, they all converge at a common regulatory node: the phosphorylation of eIF2α by one of four stress-specific kinases (HRI, PERK, GCN2, or PKR), resulting in global attenuation of protein synthesis while selectively upregulating stress-associated transcription factors, such as ATF4 [6–8]. In the context of mitochondrial dysfunction, the key molecule to trigger the ISR is DELE1 [9–11].

DELE1 localization serves as a signal for sensing and relaying mitochondrial stress. Under conditions such as mitochondrial depolarization (e.g., CCCP treatment) or ATP synthase inhibition (e.g., oligomycin treatment), DELE1 during its import is cleaved by a mitochondrial inner membrane protease, OMA1. The cleaved product of DELE1 accumulates in the cytosol, where it oligomerizes to form a scaffold platform that recruits and activates the eIF2α kinase HRI [12]. Under iron deficiency conditions, the full-length DELE1 is stabilized on the outer mitochondrial membrane (OMM) to activate HRI [11]. While the molecular mechanisms underlying ISR activation triggered by mitochondrial dysfunction are well understood, the role of DELE1-mediated ISR activation in regulating mitochondrial homeostasis remains unclear.

Mitophagy, a specialized form of autophagy that selectively eliminates damaged mitochondria, is another critical mechanism of mitochondrial quality control [13, 14].

Deficient mitophagy, leading to the accumulation of dysfunctional mitochondria, is linked to diverse human disorders, including neurodegenerative diseases such as Parkinson’s Disease (PD) [15]. This link is highlighted by mutations in two genes *PINK1* and *PRKN*, which cause familial forms of PD [16, 17]. PINK1 and PRKN play central roles in mitophagy [18]. *PINK1* encodes for PTEN-induced putative kinase 1 (PINK1), and *PRKN* encodes a E3-ubiquitin ligase, PARKIN. Under normal conditions, PINK1 is efficiently imported into healthy mitochondria. During import, PINK1 is cleaved by the matrix processing peptidase (MPP), and PARL, a mitochondrial protease located in the inner membrane [19–21]. The resulting cleaved form of PINK1 is subsequently released into the cytoplasm and degraded by the proteasome following the N-end rule [22].

However, when PINK1 import is disrupted due to mitochondrial damage, it can stabilize on the OMM, where it phosphorylates ubiquitin and PARKIN [23, 24]. As a result, PARKIN is activated and tethered on the OMM, where it further ubiquitinates OMM proteins [25–27], marking them for degradation by proteasomes and for recognition by the autophagy receptors to initialize autophagosome formation. This ultimately leads to the engulfment and degradation of the damaged mitochondria [28–30].The PINK1- PARKIN-dependent mitophagy pathway is robustly triggered by mitochondrial depolarization, but is less responsive to other non-depolarizing stress conditions, such as oligomycin treatment [24, 31].

Here, we investigated the relationship between the DELE1-HRI axis of ISR pathway and PINK1-PARKIN-dependent mitophagy under both depolarizing and non-depolarizing stress conditions. Strikingly, we found that loss of the ISR pathway components such as OMA1, DELE1, HRI, and eIF2α phosphorylation, but not ATF4, robustly activates PINK1-dependent mitophagy under non-depolarizing stress through PINK1 stabilization. ISRIB, a small molecule inhibitor of the ISR that reverses the effects of eIF2α phosphorylation [32, 33], can also enhance mitophagy, albeit to a lesser degree compared to genetic ablation of the ISR. We further showed that PINK1 stabilization is coupled with impaired mitochondrial protein import in the absence of ISR, but without mitochondrial membrane potential loss. We speculate that general protein synthesis attenuation due to the ISR activation reduces the protein influx into mitochondria, resulting in efficient mitochondrial import, even in the presence of stress. Consequently, mitophagy is inhibited under the non-depolarizing stress condition because of efficient PINK1 import and destabilization. Indeed, mildly reducing protein synthesis in ISR-deficient cells can increases mitochondrial protein import, decrease PINK1 stability and reduces mitophagy. Similarly, triggering alternative ISR pathways can suppress the mitophagy phenotype in cells without DELE1 axis of ISR, suggesting that the ISR pathways in general can regulate mitochondrial protein import and mitophagy.

## Results

### DELE1 negatively regulates mitophagy under multiple mitochondrial stress conditions

To monitor mitophagy under different cellular stress conditions, we stably expressed *mtKeima* in a HEK293T cell line equipped with CRISPRi machinery (Figure 1a). *mtKeima* encodes a mitochondrially localized red fluorescent protein with dual excitation wavelengths that vary depending on the pH of its environment. It can be excited at 440 nm, a peak predominant at pH levels above 6 as found in mitochondria, and at 586 nm, a peak predominant at pH levels below 5 as in the lysosome [34]. We measured and quantified mitophagy using flow cytometry and defined a mitophagy cell population with higher mtKeima (lysosome)/mtKeima (mitochondria) ratio (Figure 1a). CRISPRi machinery allows for the knockdown of candidate genes via expressing gene-specific sgRNAs [35]. In HEK293T cells with a non-targeting control sgRNA (*NTC*), which serves as a wild type control, we observed a very low basal level of mitophagy (about 2%). Mitochondrial stress conditions induced by oligomycin, an ATP synthase inhibitor, or carbonyl cyanide m-chlorophenyl hydrazone (CCCP), a mitochondrial uncoupler that depolarizes mitochondrial membrane potential [36], do not induce mitophagy (Supplementary Figure 1). Interestingly, knockdown of *DELE1* (*DELE1* KD) enhances basal levels of mitophagy (from 2% to 10%), which is further amplified slightly by oligomycin treatment (about 15%), but not by CCCP treatment (Supplementary Figure 1), suggesting that DELE1 may suppress mitophagy under non-depolarizing stress conditions such as oligomycin treatment.

**Figure 1.**
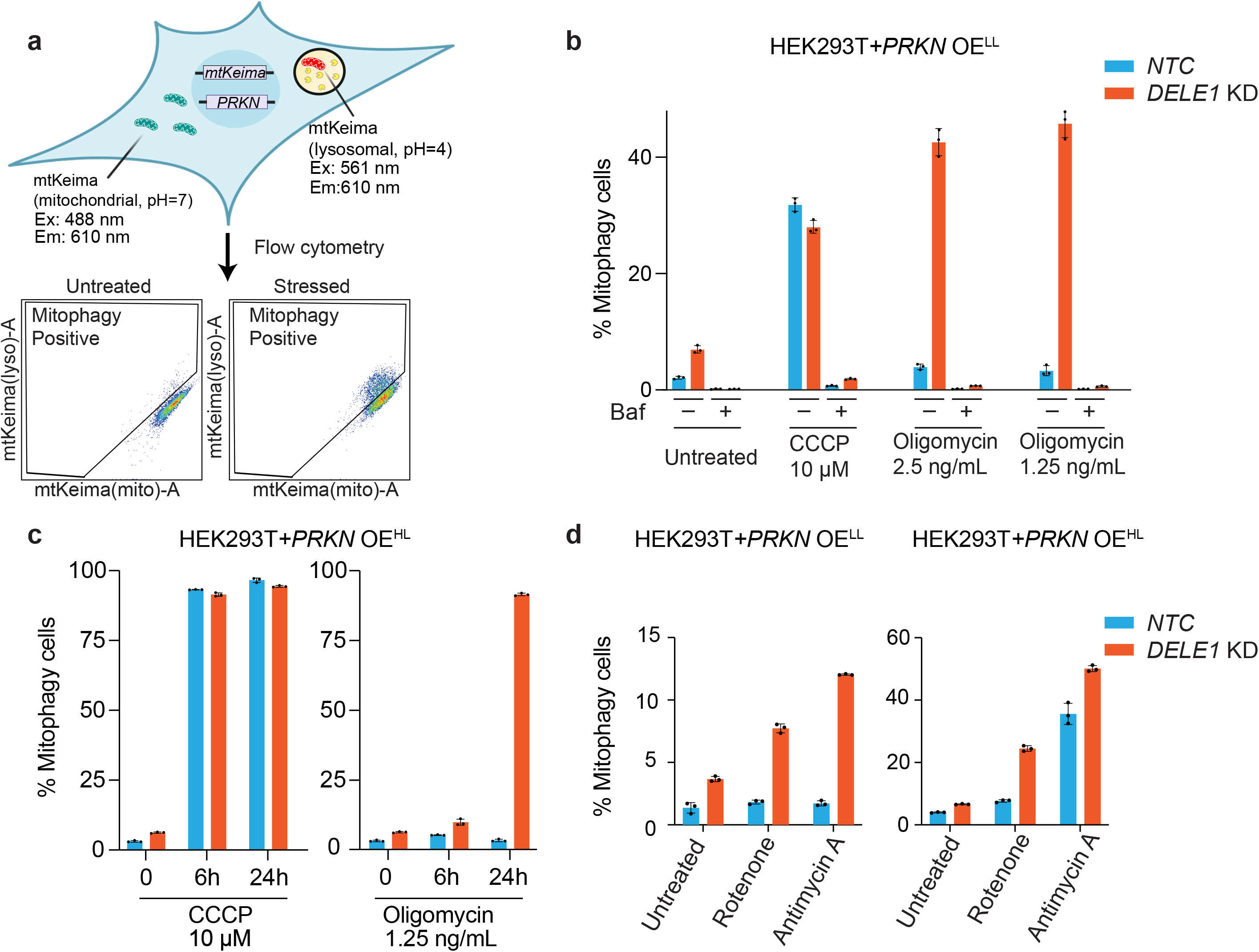
DELE1 negatively regulates mitophagy under multiple stress conditions. **(a)** Schematic illustration for monitoring mitophagy using *mtKeima* reporter via flow cytometry. Cells with increased 561 nm/405 nm mtKeima ratios compared to non-stressed cells were considered as mitophagy-positive. The diagram is generated using Biorender. **(b)** HEK293T *mtKeima* reporter cells with overexpression (OE) of *PRKN* at a lower level (*PRKN* OE^LL^) expressing a non-targeting control sgRNA (*NTC*) or sgRNA targeting *DELE1* (*DELE1* KD) were left untreated or treated with 10 μM CCCP, 1.25 ng/ml or 2.5 ng/ml oligomycin with or without 100 nM bafilomycin A (Baf) for 24 hr, followed by measurement of mitophagy using flow cytometry. (mean ± s.d., n = 3 culture wells) **(c)** HEK293T *mtKeima* reporter cells with overexpression of *PRKN* at a higher level (*PRKN* OH^LL^) expressing a non-targeting control sgRNA (*NTC*) or sgRNA targeting *DELE1* (*DELE1* KD) were left untreated or treated with 10 μM CCCP or 1.25 ng/ml oligomycin for 6 and 24 hr, followed by measurement of mitophagy using flow cytometry. (mean ± s.d., n = 3 culture wells) **(d)** HEK293T *mtKeima* reporter cells with *PRKN* OE^LL^ (left) and *PRKN* OH^HL^ (right) expressing a non-targeting control sgRNA (*NTC*) or sgRNA targeting *DELE1* (*DELE1* KD) were treated with 100 nM rotenone or 100 nM antimycin for 24 hr, followed by measurement of mitophagy using flow cytometry. (mean ± s.d., n = 3 culture wells)

We speculate that the low level of mitophagy observed in HEK293T cells could be due to a weak expression of *PRKN*. Next, we tested whether overexpression of *PRKN* would lead to a stronger phenotype. We lentivirally integrated *PRKN* into the *mtKeima* cell lines and established two cell lines with low (*PRKN* OE^LL^) and high (*PRKN* OE^HL^) *PRKN* expression, respectively. In the *PRKN* OE^LL^ cells, CCCP can induce mitophagy in about 35% of cells (Figure 1b), while in the *PRKN* OE^HL^ cells, CCCP can induce mitophagy in more than 90% of cells (Figure 1c), suggesting that the level of mitophagy induction is correlated with the *PRKN* expression levels. Both *NTC* and *DELE1* KD cells have similar levels of mitophagy following CCCP treatment (Figure 1b, c), as well as co-treatment with oligomycin and antimycin A (OA), another mitochondrial depolarizing stressor (Supplementary Figure 2). However, oligomycin only induces mitophagy in about 10% *NTC* cells regardless of *PRKN* expression level, suggesting that *PRKN* is not a limiting factor of oligomycin-induced mitophagy in wild type cells (Figure 1b, c). Strikingly, oligomycin treatment for 24 hr induces mitophagy in the absence of *DELE1* to levels comparable to CCCP treatment (Figure 1b), albeit with slower kinetics (Figure 1c). The enhanced mitophagy is significantly inhibited by Bafilomycin A, an inhibitor of autophagy that blocks the H+-ATPases (Figure 1b). In Hela cells, we observed that oligomycin can also induce significantly higher levels of mitophagy in *DELE1* KD cells, although this effect requires higher concentrations (Supplementary Figure 3). Additionally, the levels of oligomycin-induced mitophagy are not as high as those triggered by CCCP. These results suggest that oligomycin sensitivity varies across different cell lines.

In addition to oligomycin, we observed increased mitophagy in *DELE1* KD cells under other non-depolarizing stress conditions such oxidative phosphorylation (OXPHOS) Complex I inhibitor rotenone and Complex III inhibitor antimycin A (Figure 1d) in a PARKIN-level dependent manner. Notably, oligomycin treatment triggers the strongest mitophagy in *DELE1* KD cells among all OXPHOS inhibitors, prompting us to use oligomycin as the primary stressor in our subsequent mechanistic studies.

### Oligomycin induces PINK1 stabilization on the mitochondria in cells without DELE1

Given that mitophagy induction in *DELE1* KD cells is much stronger in *PRKN* OE cells (Figure 1b, c), it is likely that oligomycin-induced mitophagy in *DELE1* KD cells occurs through the PINK1-PARKIN pathway. To test this, we established *PINK1* knockout (KO) cell lines and measured mitophagy. Similar to CCCP treatment, oligomycin-induced mitophagy in *DELE1* KD cells is abolished in the absence of PINK1 (Figure 2a). PINK1 trafficking is coupled with its stability. Under normal conditions, PINK1 is imported into mitochondria and cleaved by PARL [19–21]. The truncated PINK1 then relocates to the cytoplasm where it is degraded following the N-end rule [22]. Upon mitochondrial membrane depolarization, PINK1’s import is compromised, and it is stabilized on the OMM, where PINK1 can recruit the E3 ubiquitin ligase PARKIN to label the damaged mitochondria for lysosomal degradation [18]. To test whether PINK1 is stabilized following oligomycin treatment in the absence of DELE1, we immunoblotted PINK1 in both wild type and *DELE1* KD cells under CCCP and oligomycin conditions. Consistent with previous studies, CCCP induces strong stabilization of full length PINK1. While oligomycin treatment for 24 hr does not stabilize PINK1 in wild type cells, PINK1 is significantly accumulated in *DELE1* KD cells (Figure 2b). We further performed subcellular fractionation and detected that PINK1 accumulates in the mitochondrial fraction in *DELE1* KD cells under oligomycin condition (Figure 2c). In addition, we immunoblotted the autophagy marker LC3, and found that LC3-II, a lipidated form that is associated with autophagosome levels [37], is significantly higher in *DELE1* KD cells after oligomycin treatment, indicating a higher level of autophagy. Lastly, we immunoblotted the mitochondrial membrane protein COXIV, whose level is used as an indicator of mitophagy. We found that oligomycin treatment significantly reduces the level of COXIV in *DELE1* KD cells, indicating a high level of mitophagy (Figure 2b).

**Figure 2.**
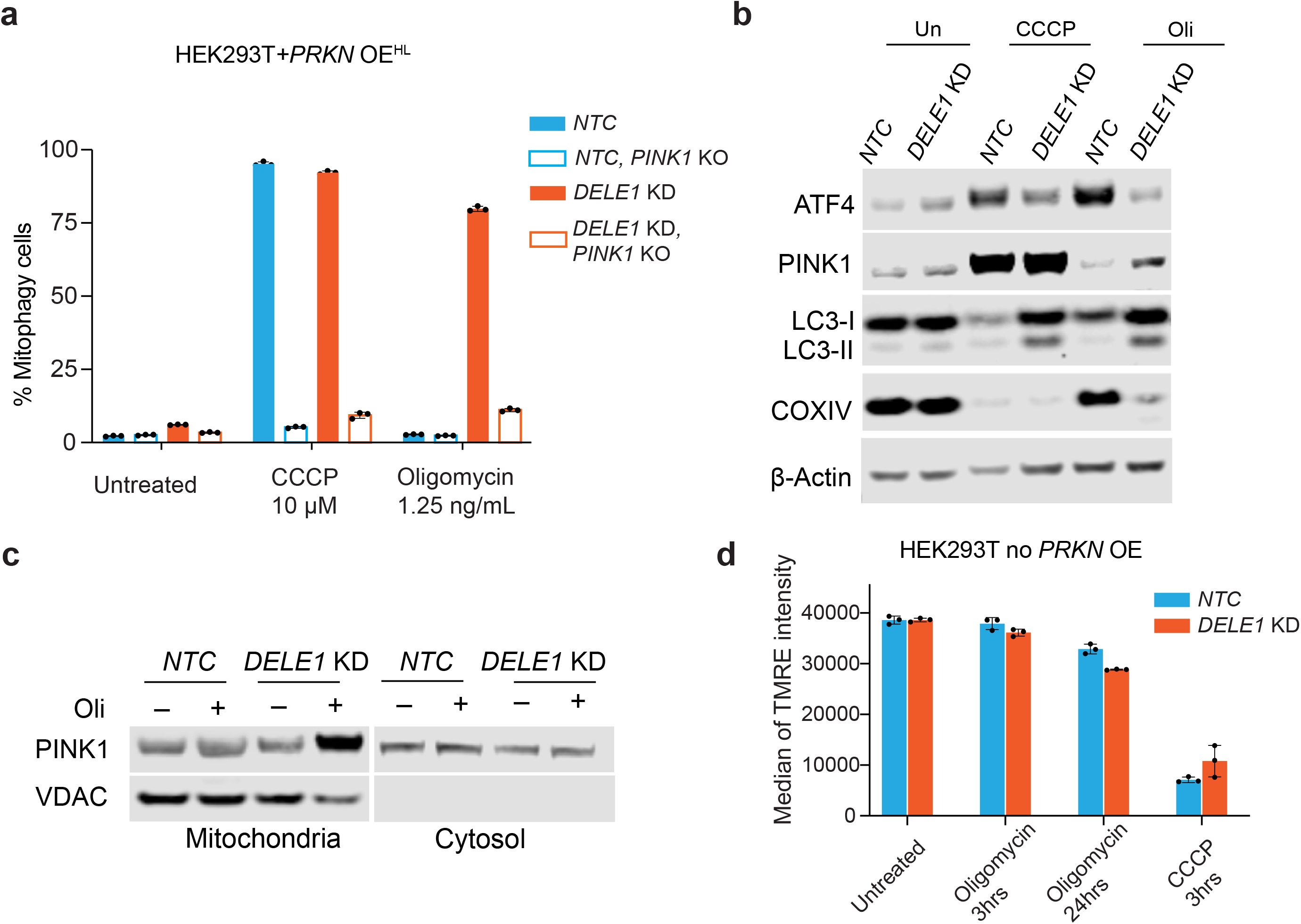
Oligomycin promotes PINK1 accumulation on mitochondria in DELE1- knockdown cells. **(a)** HEK293T *PRKN* OE^HL^ wild type (WT) or *PINK1* (KO) cells with or without *DELE1* KD were treated with 10 μM CCCP or 1.25 ng/ml oligomycin for 24 hr, followed by measurement of mitophagy using flow cytometry. (mean ± s.d., n = 3 culture wells) **(b)** Immunoblots of ATF4, PINK1, LC3 and COXIV. HEK293T *PRKN* OE^HL^ cells expressing a non-targeting control sgRNA (*NTC*) or sgRNA targeting *DELE1* (*DELE1* KD) were treated with 10 μM CCCP or 1.25 ng/mL of oligomycin for 24 hr. β-actin was used as the loading control. **(c)** Immunoblots of PINK1 and VDAC. Mitochondrial and cytosol fractions were isolated from HEK293T cells expressing a *NTC* or *DELE1* sgRNAs. Cells were treated with 1.25 ng/mL of oligomycin for 24 hr before mitochondrial isolation. **(d)** HEK293T cells with a *NTC* or *DELE1* sgRNAs were treated with 10 μM CCCP for 3 hr, as a positive control for mitochondrial depolarization, and 1.25 ng/ml oligomycin for 3 for 24 hr, followed by 100 nM TMRE staining and flow cytometry analysis. (mean ± s.d., n = 3 culture wells)

Notably, even though the PINK1 level in *DELE1* KD cells following oligomycin treatment is significantly lower than those following CCCP treatment in both wild type and *DELE1* KD cells, it still reaches a medium threshold level to trigger a similar level of mitophagy (Figure 2b and Figure 1b, c), but with slower kinetics (Figure 1c) [38]. These results, mirroring the mtKeima analysis, suggest that DELE1 negatively regulates mitophagy by modulating PINK1 stability.

To understand the mechanism of PINK1 stabilization, we measured the mitochondrial membrane potential, as loss of mitochondrial membrane potential is a known mechanism that stabilizes PINK1 and induces mitophagy [24]. Previous studies suggested that oligomycin can hyperpolarize mitochondria during the initial stage and eventually slightly depolarize the mitochondria [39]. Here, we detected oligomycin treatment for 3 hr does not change mitochondrial membrane potential and only slightly reduced the membrane potential after 24 hr treatment in wild type cells (Figure 2d). We did notice a small reduction of TMRE signal in *DELE1* KD cells following oligomycin treatment compared to wild type cells, even though it is still significantly higher than the membrane potential after depolarization by CCCP (Figure 2d). We speculate this slight reduction maybe due to a smaller number of mitochondria, a result of slightly higher mitophagy in *DELE1* KD cells (Supplementary Figure 1). These results suggest that a different mechanism, independent of mitochondrial membrane potential, regulates PINK1 stabilization in the absence of DELE1 following oligomycin treatment.

### The absence of DELE1 reduces mitochondrial protein import efficiency

Previous studies have shown that impairment of mitochondrial import machinery can stabilize PINK1 without loss of mitochondrial membrane potential [31, 40]. Therefore, we hypothesize that DELE1 pathway may play a role in maintaining mitochondrial protein import under non-depolarizing stress conditions such as oligomycin used here. To investigate this, we conducted three different assays to monitor mitochondrial protein import.

First, we performed immunoblotting for HSPD1, a mitochondrial matrix resident protein. HSPD1 is translated as a precursor form containing a mitochondrial targeting sequence (MTS), which is cleaved upon successful import into mitochondria. The accumulation of the precursor form serves as an indicator of impaired mitochondrial import [31]. As expected, we observed a slight accumulation of HSPD1 precursor, indicating import failure, following CCCP treatment, but not oligomycin treatment in wild type cells (Figure 3a). Notably, *DELE1* KD cells exhibited a stronger accumulation of the HSPD1 precursor form, after both CCCP and oligomycin treatments (Figure 3a). These results suggest that DELE1 positively regulates mitochondrial import under both stress conditions. Consistently, we observed a further accumulation of PINK1 even under CCCP treatment in *DELE1* KD cells compared to *WT* cells (Figure 2b). However, this increase in PINK1 level does not further enhance mitophagy, as CCCP treatment in *WT* cells already results in sufficient PINK1 accumulation to trigger mitophagy.

**Figure 3.**
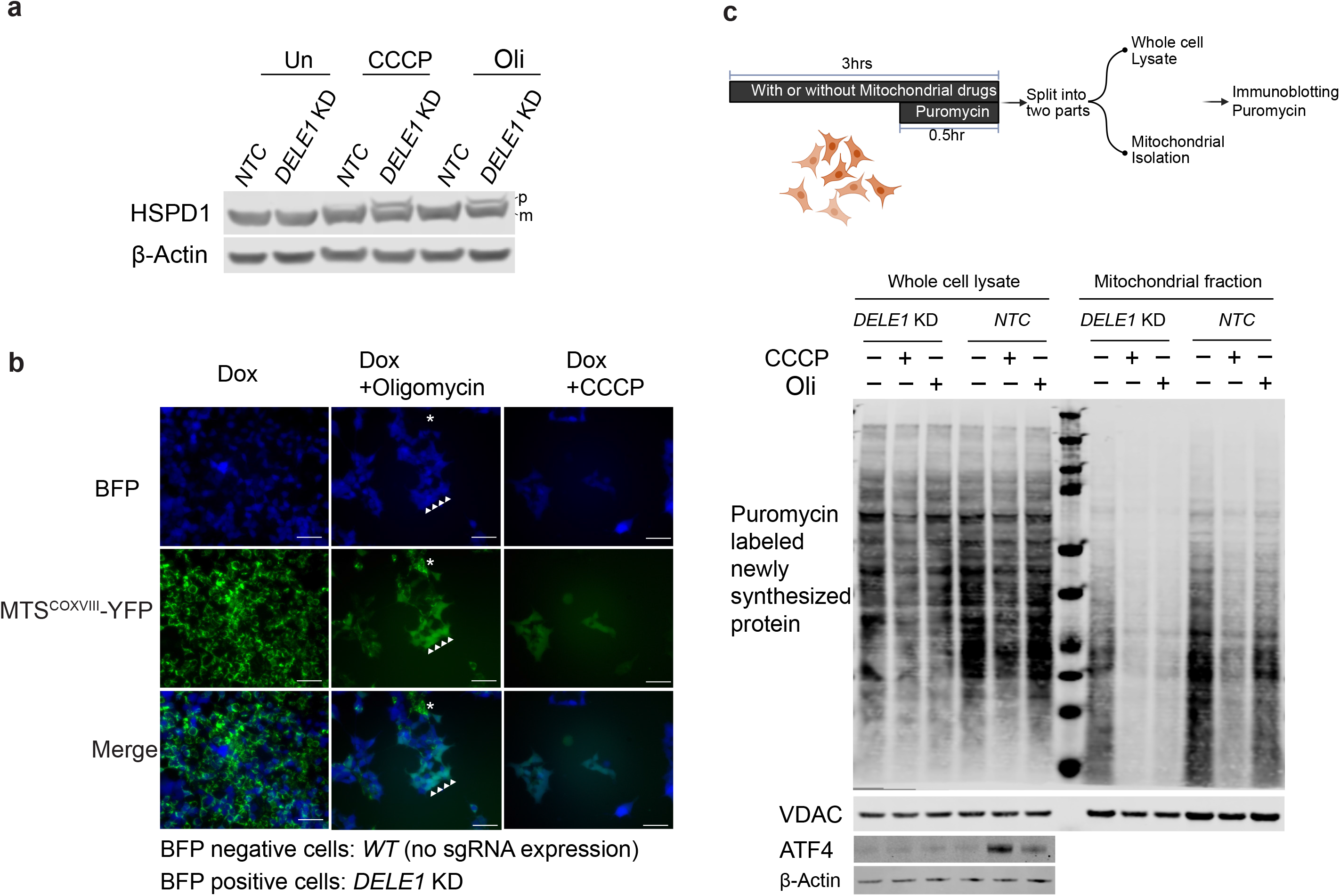
DELE1 maintains mitochondrial protein import upon stress. **(a)** Immunoblots of HSPD1. HEK293T cells with *NTC* or *DELE1* sgRNAs (*DELE1 KD*) were treated with 10 μM CCCP or 1.25 ng/mL oligomycin (Oli) for 24 hr. p: precursor form of HSPD1. m: mature form of HSPD1. β-actin was used as a loading control. **(b)** Inducible *MTS^COXVIII^-YFP* cells were infected with *DELE1* sgRNAs to generate a mixed population with about 50% of cells containing sgRNA (BFP+) and remaining cells without BFP (BFP-) serving as wild type (*WT*) control. These cells were treated with 500 ng/mL doxycycline with or without 10 μM CCCP or 1.25 ng/mL oligomycin for 24 hr. Asterisk highlights some BFP negative cells as wild type control. Arrowheads highlight some BFP positive cells (*DELE1* KD). Scale bar 100 μm. **(c)** Top: Experimental scheme of puromycin labeling assay to evaluate total mitochondrial protein import efficiency (generated using Biorender). Cells were left untreated or treated with mitochondrial drugs for 3 hr. Puromycin was added during the last 30 min of drug treatments to label the newly synthesized protein. Mitochondria were isolated for immunoblotting. Bottom: Immunoblot of puromycin in isolated mitochondria and whole cell lysate. Cells with *NTC* or *DELE1* sgRNAs were treated with 10 μM CCCP and 1.25 ng/mL oligomycin for 3 hr. ATF4 activation indicates mitochondrial stress and *DELE1* knockdown. β-Actin (whole cell) and VDAC (mitochondrial fraction) serve as the loading control.

Second, to rule out the possibility that the HSPD1 precursor accumulation in *DELE1* KD cells after oligomycin treatment was due to higher protein translation (a consequence of the ISR inactivation) but not due to a reduction of mitochondrial import efficiency, we made an inducible reporter containing a mitochondrial targeting sequence from *COXVIII* fused to *YFP*. We induced the expression of *MTS^COXVIII^-YFP* concurrently with oligomycin treatment and observed that MTS^COXVIII^-YFP maintained mitochondrial localization in wild type cells (Figure 3b, Supplementary Figure 4a). In contrast, in a significant population of *DELE1* KD cells, MTS^COXVIII^-YFP lost their mitochondrial localization (Figure 3b, Supplementary Figure 4a), suggesting that mitochondrial import of the reporter was impaired.

Finally, we combined puromycin labeling of newly synthesized proteins with subcellular fractionation experiments to measure the import efficiency across a wide range of proteins (Figure 3c). Puromycin, which mimics aminoacyl-tRNAs, is incorporated into proteins during translation [41]. We added puromycin during the final 30 minutes of drug treatment and used immunoblotting to detect puromycin-labeled proteins in isolated mitochondria as a proximity for overall mitochondrial import efficiency. Strikingly, *DELE1* KD cells displayed a significantly reduced amount of newly synthesized proteins in the mitochondria following oligomycin treatment, indicating that the mitochondrial import deficiency in these cells extends beyond specific proteins like HSPD1 or MTS^COXVIII^-YFP, affecting a broader range of mitochondrial substrates (Figure 3c).

In summary, our findings suggest that DELE1 plays a crucial role in maintaining mitochondrial protein import under stress conditions.

### The ISR pathway inhibits mitophagy by maintaining mitochondrial protein import via reduced protein synthesis

Mitochondrial stress conditions, including both CCCP and oligomycin, trigger the OMA1- DELE1-HRI axis of ISR, resulting in reduced protein synthesis and increased ATF4 translation [9, 10]. Next, we set out to determine if DELE1 maintains mitochondrial import and inhibits mitophagy through ATF4 upregulation downstream of OMA1, HRI and eIF2α phosphorylation. Similar to *DELE1* knockdown, loss of OMA1, which cleaves DELE1 upon oligomycin treatment, significantly enhances oligomycin-induced mitophagy (Figure 4a). Additionally, *HRI* knockdown (Figure 4a) cells and cells deficient in eIF2α phosphorylation (*eIF2α^S49/52/A^*) (Figure 4c, supplementary Figure 7) both exhibit strong mitophagy upon oligomycin treatment. At the same time, we observed PINK1 accumulation and mitochondrial import impairment in *HRI* KD (Figure 4b, Supplementary Figure 4a, b) and *eIF2α^S49/52/A^* cells (Figure 4d, Supplementary Figure 4a) upon oligomycin treatment. In contrast, *ATF4* knockout cells do not induce mitophagy (Figure 4e) or impair mitochondrial protein import (Figure 4f, Supplementary Figure 4a) following oligomycin treatment. Additionally, measurements of mitochondrial membrane potential in cells lacking different DELE1-ISR components revealed no significant loss of membrane potential following oligomycin treatment (Supplementary Figure 4c). These results suggest that the OMA1-DELE1-HRI-eIF2α phosphorylation regulates mitochondrial protein import efficiency and mitophagy independently of ATF4 upregulation, likely by attenuating general protein synthesis.

**Figure 4.**
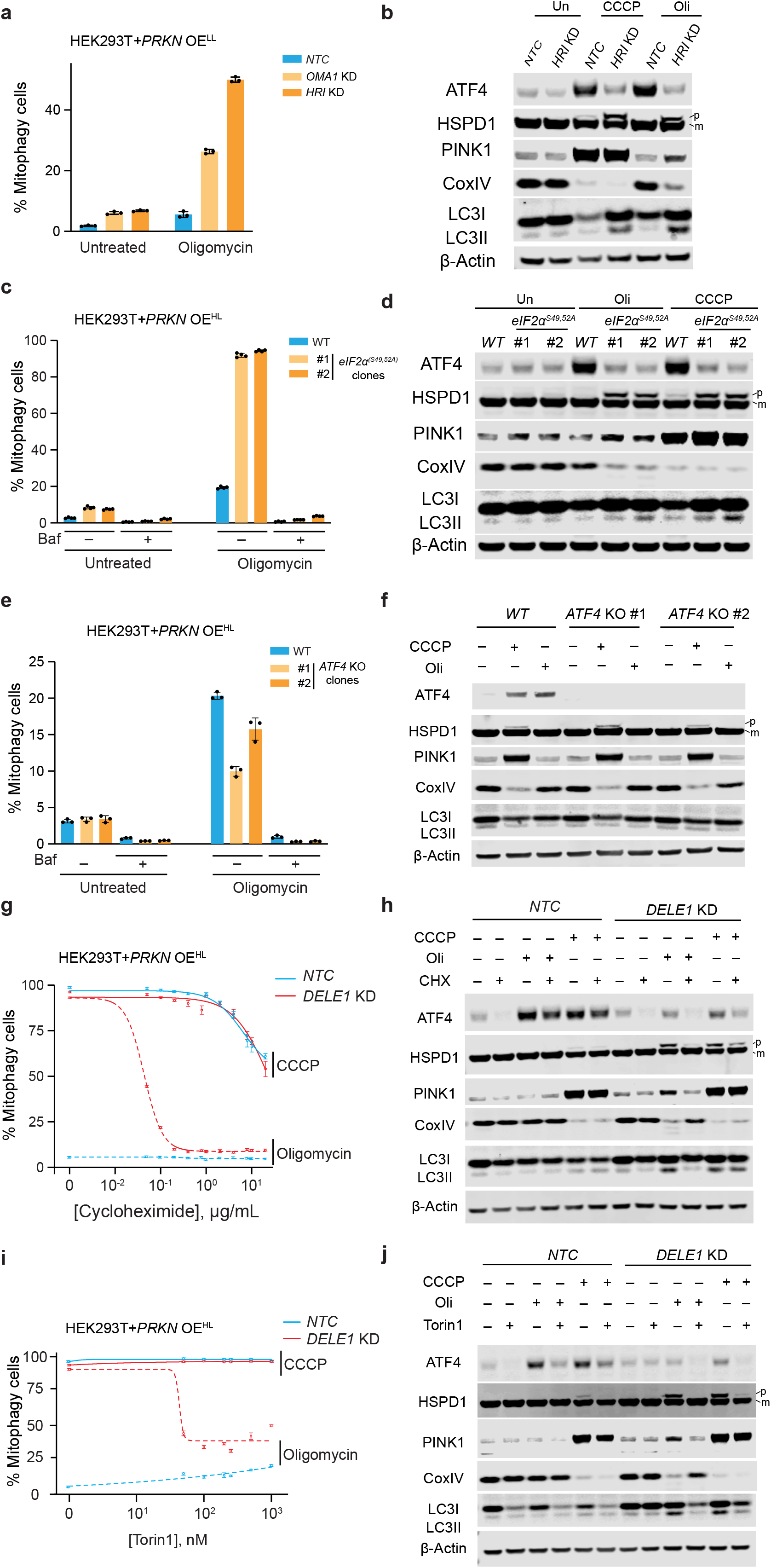
DELE1-ISR pathway negatively regulates mitophagy via attenuating protein synthesis. **(a)** HEK293T *PRKN* OE^LL^ cells expressing non-targeting control sgRNA (*NTC*), sgRNA targeting *OMA1* (*OMA1* KD) or *HRI* (*HRI* KD) were left untreated or treated with 1.25 ng/mL oligomycin for 24 hr, followed by measurement of mitophagy via flow cytometry. (mean ± s.d., n = 3 culture wells) **(b)** Immunoblots of ATF4, HSPD1, PINK1, CoxIV, LC3, and β-Actin. *NTC* and *HRI* KD cells were left untreated or treated with 10 μM CCCP or 1.25 ng/mL oligomycin for 24 hr. β-Actin serves as loading control. **(c)** Wild type (*WT*) or two clonal *eIF2α^S49/52/A^* cells with *PRKN OE^HL^*were left untreated or treated with 1.25 ng/mL oligomycin with or without bafilomycin A (Baf), followed by measurement of mitophagy via flow cytometry. (mean ± s.d., n = 3 culture wells) **(d)** Immunoblots of ATF4, HSPD1, PINK1, CoxIV, LC3, and β-Actin in *WT* and *eIF2α^S49/52/A^* cells following 1.25 ng/mL oligomycin for 24 hr. β-Actin serves as the loading control. **(e)** Flow cytometry measurement of mitophagy in *WT* or two *ATF4* KO clonal cell lines with *PRKN* OE^HL^ following treatment with 1.25 ng/mL oligomycin for 24 hr in the presence or absence of bafilomycin A (Baf). (mean ± s.d., n = 3 culture wells) **(f)** Immunoblots of ATF4, HSPD1, PINK1, CoxIV and LC3 and β-Actin in WT and two *ATF4* KO clonal cell lines following 10 μM CCCP or 1.25 ng/mL oligomycin treatment for 24 hr. β-Actin serves as the loading control. **(g)** *NTC* and *DELE1* KD cells with *PRKN OE^HL^*were treated with 10 μM CCCP or 1.25 ng/mL oligomycin in the presence of cycloheximide at 12 different concentrations (0, 50 ng/mL, 100 ng/mL, 200 ng/mL, 400 ng/mL, 800 ng/mL, 1 μg/mL, 2 μg/mL, 4 μg/mL, 8 μg/mL,10 μg/mL and 20 μg/mL) for 24 hr followed by flow cytometry to measure mitophagy. Cycloheximide concentrations were converted to their base-10 logarithmic values. A nonlinear regression analysis using a log(inhibitor) vs. response model with a variable slope (four parameters) was performed to generate the plot. (mean ± s.d., n = 3 culture wells) **(h)** Immunoblots of ATF4, HSPD1, PINK1, CoxIV and LC3 and β-Actin in *WT* and *DELE1* KD cells with *PRKN OE^HL^* following 10 μM CCCP or 1.25 ng/mL oligomycin treatment for 24 hr with or without 100 ng/mL cycloheximide (CHX). p: precursor; m: mature. β-Actin serves as the loading control. **(i)** *NTC* and *DELE1* KD cells with *PRKN* OE^HL^ are treated with 10 μM CCCP or 1.25 ng/mL oligomycin in the presence of torin1 at 7 different concentrations (0, 50, 100, 200, 250, 500 and 1000 nM) for 24 hr followed by flow cytometry to measure mitophagy. Torin1 concentrations were converted to their base-10 logarithmic values. A nonlinear regression analysis using a log(inhibitor) vs. response model with a variable slope (four parameters) was performed to generate the plot. (mean ± s.d., n = 3 culture wells) **(j)** Immunoblots of ATF4, HSPD1, PINK1, CoxIV and LC3 and β-Actin in NTC and *DELE1* KD cells with *PRKN* OE^HL^ following 10 μM CCCP or 1.25 ng/mL oligomycin treatment for 24 hr with or without 250 nM torin1. p: precursor; m: mature. β-Actin serves as the loading control.

Overexpression of mitochondrial proteins, particularly those harboring a bipartite signal, can cause mitochondrial import defects in yeast and mammals [42, 43]. We hypothesize that, in the absence of the ISR pathway, because protein synthesis rate is not attenuated, the amount of mitochondrial protein precursors synthesized becomes overwhelming for stressed mitochondria, causing mitochondrial import deficiency. This import deficiency, in turn, stabilizes PINK1 and enhances mitophagy. To test this hypothesis, we treated cells with the translation inhibitor cycloheximide to reduce protein synthesis and examined its effects on mitophagy in ISR-deficient cells.

Both wild type and *DELE1* KD cells were treated with varying concentrations of cycloheximide concurrently with CCCP or oligomycin. For these experiments, we selected sub-optimal concentrations of cycloheximide compared to those typically used in pulse-chase experiments, given that PINK1 stabilization requires ongoing protein translation. As expected, high doses of cycloheximide inhibited mitophagy in both wild type and *DELE1* KD cells treated with CCCP (Figure 4g), likely due to the strong suppression of protein synthesis, limiting PINK1 availability. However, oligomycin-induced mitophagy in *DELE1* KD cells showed significantly higher sensitivity to cycloheximide compared to CCCP-induced mitophagy. The IC_50_ of cycloheximide required for mitophagy inhibition in *DELE1* KD cells under oligomycin treatment was 0.04 μg/mL, approximately 575 times lower than the 23 μg/mL under CCCP treatment. Such a drastic difference in cycloheximide sensitivity suggests that the inhibition of oligomycin-induced mitophagy in *DELE1* KD cells is not due to a reduction in PINK1 translation. Instead, it indicates that mitochondrial protein import is restored, as evidenced by a notable reduction in the precursor form of HSPD1 in *DELE1* KD cells, but not in wild-type cells under mitochondrial stress conditions, leading to PINK1 destabilization (Figure 4h).

Next, we investigated whether mTOR inhibition, which also attenuates protein translation, could suppress mitophagy in ISR-deficient cells. Wild type and *DELE1* KD cells were co-treated with Torin1, a potent mTOR inhibitor, alongside oligomycin or CCCP. Although Torin1 by itself promotes general autophagy [44] and hence slightly enhances mitophagy (Supplementary Figure 5), it significantly reduces oligomycin-induced mitophagy in *DELE1* KD cells, but has no effect on CCCP-induced mitophagy, as shown by the *mtKeima* reporter assays (Figure 4i). Consistent with this, immunoblotting results showed elevated CoxIV levels in *DELE1* KD cells co-treated with oligomycin and Torin1, along with PINK1 destabilization, possibly due to improved mitochondrial import efficiency, as suggested by the reduction in the precursor form of HSPD1 (Figure 4j). Interestingly, unlike cycloheximide treatment, Torin1 co-treatment with CCCP also resulted in a notable reduction of the HSPD1 precursor form and PINK1 levels in wild-type cells. This reduction is unlikely to result from translation inhibition, as neither HSPD1 nor PINK1 have been identified as mTOR targets [45]. These results suggest mTOR inhibition via Torin1 improve mitochondrial import even under depolarizing stress. We also observed a reduction in ATF4 following Torin1 treatment (Figure 4j), in line with previous reports that mTOR activity is required for ATF4 translation, independent of eIF2α phosphorylation [46]. However, since ATF4 does not regulate mitophagy or mitochondrial import (Figure 4e, f), Torin1 and ISR may have an additive effect on protein translation, improving mitochondrial import under CCCP condition in wild type cells. Although PINK1 levels are reduced in wild type or *DELE1* KD cells following co-treatment with CCCP and Torin1, they remain sufficient to trigger mitophagy (Figure 4i, j).

These findings indicate that non-depolarizing stress, such as oligomycin treatment, increases the sensitivity of the mitochondrial import machinery to protein synthesis rates. When the ISR is inhibited, the mitochondrial import machinery becomes overwhelmed by mitochondrial substrates. Attenuating protein synthesis, either through a general translation inhibition or mTOR inhibition, can alleviate mitochondrial import stress and suppress mitophagy. In contrast, under depolarizing stress, the mitochondrial import machinery is more severely compromised and less responsive to changes in protein synthesis rates.

### Modulating ISR using small molecules can regulate mitophagy

We next sought to investigate whether modulating the integrated stress response (ISR) through small-molecules inhibition or activation could regulate mitophagy. ISRIB, an ISR inhibitor that antagonizes phosphorylated eIF2α, promotes general protein translation while inhibiting ATF4 translation [32, 33]. To investigate the effect of ISRIB on mitophagy induced by oligomycin, wild type cells were treated with ISRIB at concentrations ranging from 10 nM to 2000 nM. Similar to *DELE1* knockdown, ISRIB significantly promotes mitophagy following oligomycin treatment (Figure 5a). However, the extent of mitophagy induction by ISRIB was less pronounced compared to *DELE1* knockdown. This difference may be due to ISRIB not fully blocking the ISR, as suggested by ATF4 induction (Figure 5b). As a result, only a small accumulation of HSPD1 precursor and PINK1 were observed (Figure 5b). Interestingly, ISRIB exerts a stronger effect on CCCP treatment than oligomycin treatment, suggesting that unknown mechanisms associated with oligomycin may counteract the action of ISRIB.

**Figure 5.**
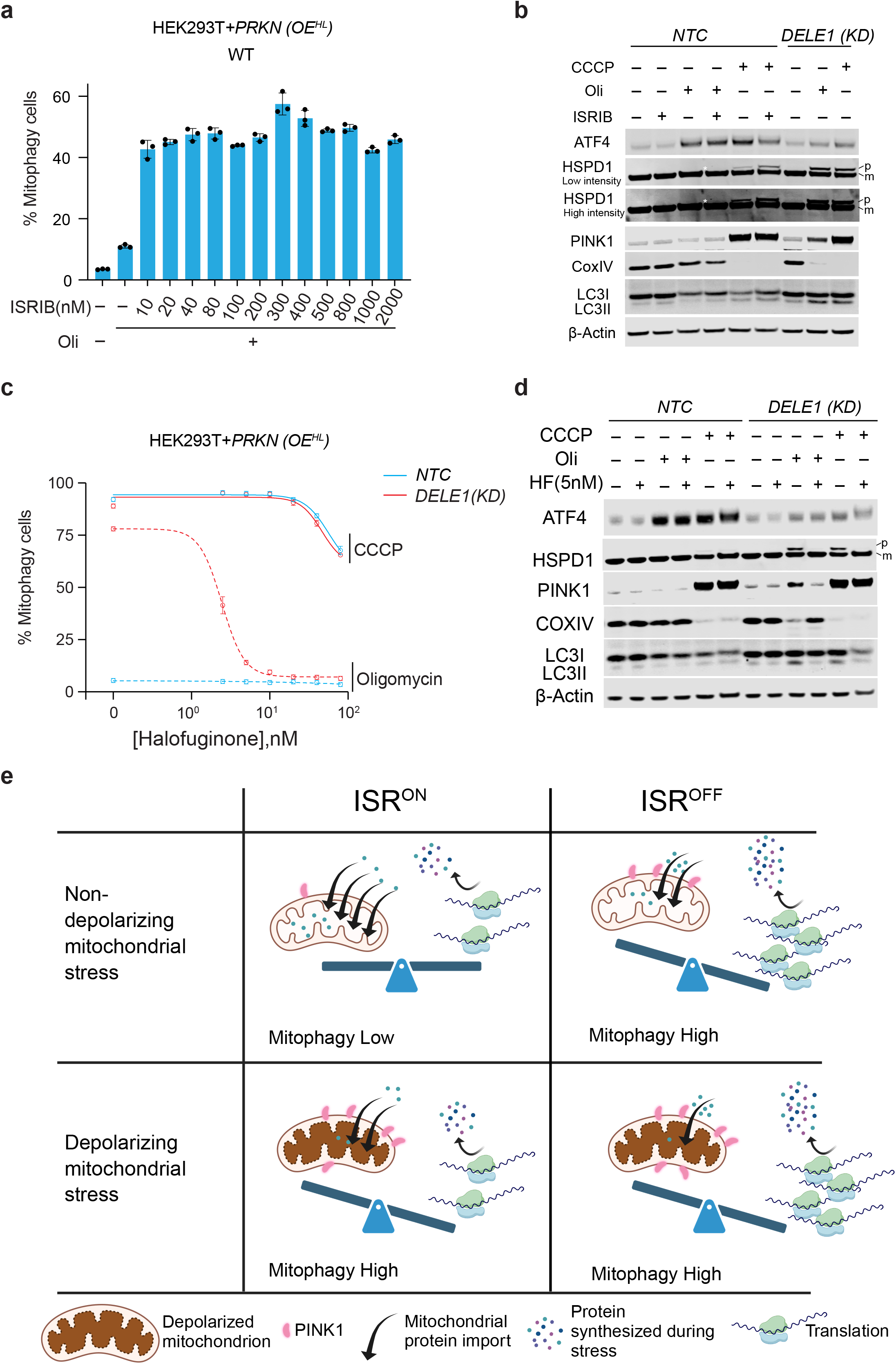
Modulation of mitophagy by small molecules that inhibit or activate the ISR. **(a)** Measurement of mitophagy in wild type (*WT*) HEK293T with *PRKN* OE^HL^ treated with 1.25 ng/mL oligomycin for 24 hr with ISRIB at 13 different concentrations. (mean ± s.d., n = 3 culture wells) **(b)** Immunoblots of ATF4, HSPD1, PINK1, CoxIV and LC3 and β-Actin in *NTC* and *DELE1* KD cells with *PRKN* OE^HL^ following 10 μM CCCP or 1.25 ng/mL oligomycin treatment for 24 hr with or without 300 nM ISRIB. p: precursor; m: mature. β-Actin serves as the loading control. **(c)** *NTC* and *DELE1 KD* cells with *PRKN* OE^HL^ were treated with 10 μM CCCP or 1.25 ng/mL oligomycin in the presence of halofuginone at 7 different concentrations (0, 2.5 nM, 5 nM, 10 nM, 20 nM, 40 nM and 80 nM) for 24 hr followed by flow cytometry to measure mitophagy. Halofuginone concentrations were converted to their base-10 logarithmic values. A nonlinear regression analysis using a log(inhibitor) vs. response model with a variable slope (four parameters) was performed to generate the plot. (mean ± s.d., n = 3 culture wells) **(d)** Immunoblots of ATF4, HSPD1, PINK1, CoxIV and LC3 and β-Actin in *NTC* and *DELE1* KD cells with *PRKN* OE^HL^ following 10 μM CCCP or 1.25 ng/mL oligomycin treatment for 24 hr with or without 5 nM halofuginone (HF). p: precursor; m: mature. β- Actin serves as the loading control. **(e)** Model for how the ISR pathway regulates mitochondrial protein import, PINK1 stability and mitophagy under both depolarizing and non-depolarizing stress. This is generated using Biorender.

Next, we explored whether activating the ISR through alternative pathways could bypass the need for DELE1 in regulating mitochondrial import and mitophagy.

Pharmacological activation of compensatory ISR kinases has been shown to mitigate cellular and mitochondrial dysfunction resulting from impaired PERK signaling [47].

Halofuginone induces the accumulation of uncharged proline tRNA, thereby activating the GCN2 branch of the ISR [48]. Strikingly, halofuginone significantly suppresses mitophagy in *DELE1*-deficient cells treated with oligomycin, and this effect occurred at much lower concentrations than required for CCCP treatment (Figure 5c). Co-treatment with 5nM halofuginone and oligomycin significantly reduces the accumulation of the precursor form of HSPD1 and PINK1, indicating an enhanced mitochondrial import facilitated by halofuginone. This improved import leads to PINK1 destabilization and reduced mitophagy, as evidenced by the elevated levels of COXIV (Figure 5d).

Similarly, thapsigargin, a non-competitive inhibitor of the endoplasmic reticulum (ER) Ca^2+^ ATPase (SERCA) [49], which triggers ER stress and activates the PERK-mediated ISR pathway [50], also inhibits mitophagy in DELE1-deficient cells following oligomycin treatment (Supplementary Figure 6). However, tharpsigargin treatment at higher concentrations (> 20 nM) exhibits high cytotoxicity, which may contribute to low mitophagy under both oligomycin and CCCP conditions at these concentrations.

These results indicate that protein attenuation through alternative ISR pathways can preserve mitochondrial import efficiency under oligomycin treatment, which leads to PINK1 destabilization and mitophagy suppression (Figure 5e).

## Discussion

Our studies uncover a novel link between the integration stress response (ISR) activation and mitophagy under different mitochondrial stress conditions. The DELE1- HRI axis of ISR is robustly triggered in response to nearly all mitochondrial stress conditions observed in many cell types, as well as in several mitochondrial myopathy mouse models [51–54]. In contrast, while PINK1-PRKN-dependent mitophagy is strongly activated by mitochondrial depolarization induced by CCCP, it occurs at significantly lower levels under non-depolarizing stress conditions, such as those induced by oligomycin. Through our investigation of ISR activation in relation to mitophagy induction, we identified a previously unrecognized role for ISR in negatively regulating PINK1-PARKIN-dependent mitophagy.

Recently, two additional independent studies have uncovered a connection between the DELE1-HRI pathway and mitophagy, albeit with conflicting conclusions [55, 56]. Chakrabarty et al., using whole-genome screening in K562 cells, identified a positive role for the DELE1-HRI pathway in mitophagy induced by iron deficiency [55]. Their findings suggest that the HRI branch of ISR activation repurposes eIF2α phosphorylation from regulating translational initiation to initiating mitophagy in response to mitochondrial dysfunction. Furthermore, they concluded that HRI is broadly essential for mitophagy triggered by mitochondrial depolarization as well as hypoxia, as demonstrated in HeLa cells. Conversely, Singh, Agarwal, Volpi et al., found that the DELE1-HRI pathway negatively regulates mitophagy [56]. Their study began with a genetic siRNA screen targeting all known human Ser/Thr kinases in HeLa cells, revealing that *HRI* knockdown increases the stabilization and activation of PINK1, as evidenced by enhanced ubiquitin Ser65 phosphorylation following mitochondrial depolarization. Additionally, they observed that inhibiting the ISR pathway, either by genetic knockdown of *DELE1* or *HRI*, or by using the small-molecule inhibitor ISRIB, amplifies PINK1-dependent mitophagy while having little effect on iron-deficiency induced mitophagy. These contrasting findings suggest that ISR’s role in mitophagy may vary depending on cell type and the specific mitochondrial stress encountered.

Our studies provide strong evidence supporting a negative regulatory role for the ISR in mitophagy. We found that under a wide range of mitochondrial stress conditions that do not induce mitochondrial depolarization, PINK1-dependent mitophagy is significantly suppressed due to activation of the DELE1-HRI-mediated ISR. However, under depolarizing stress conditions, such as those induced by CCCP and a combination of oligomycin and Antimycin A (OA, as used in the aforementioned study [56] Supplementary Figure 2), the absence of ISR does not further enhance mitophagy. This is likely because mitophagy levels have already reached their maximum in the presence of ISR, despite the observed increase in PINK1 stabilization in the absence of ISR. This negative regulation on mitophagy by the ISR might be linked to its role in preserving mitochondrial protein import efficiency.

Mitochondrial protein import efficiency is intricately coupled to the regulation of mitochondrial homeostasis [57, 58]. Although various stressors disrupt mitochondrial function through distinct mechanisms, they generally affect mitochondrial protein import efficiency to different extents. Among mitochondrial proteins, DELE1 appears particularly sensitive to import defects due to its unusually long mitochondrial targeting sequence [59], allowing it to sense a wide range of stress conditions and trigger the ISR [9, 10, 60]. In *C. elegans*, ATFS-1 contains a less efficient MTS, enabling it to sense mitochondrial perturbations as well as mitochondrial biogenesis during development [61]. One the other hand, PINK1 import is primarily impacted by more severe import defects, such as those caused by mitochondrial depolarization [22, 23]. Recent research has shown that iron chelation halts DELE1 import, stabilizing full-length DELE1 on the outer mitochondrial membrane. Interestingly, this stress does not affect PINK1 import and stabilization [11]. Our findings here demonstrate that mitochondrial import can be positively modulated by ISR activation under various pharmacological perturbations, as well as in response to mitochondrial DNA breaks, as shown in studies by Fu et al [62].

The ISR regulates mitochondrial protein import in an ATF4-independent manner, but it relies on the rate of protein synthesis. When the ISR is inactivated, increased protein translation leads to elevated levels of mitochondrial precursor proteins, which can overwhelm the mitochondria import machinery of stressed mitochondria and exacerbate protein import defects. Indeed, we observed significant accumulation of mitochondrial precursors and a marked reduction in proteins that are imported into mitochondria in cells lacking the ISR pathway. A recent CRISPR screening study identified that the loss of genes involved in mitochondrial import machinery induces PINK1-dependent mitophagy, even without the loss of mitochondrial membrane potential [31]. Consistent with this, our study demonstrated that PINK1 becomes stabilized on the outer mitochondrial membrane to trigger mitophagy in mitochondrial-import-deficient cells due to an inactivation of the ISR following the treatment with various non-depolarizing mitochondrial drugs. Although we did not observe further enhancement of mitophagy following CCCP treatment, we did observe a further diminishment of mitochondrial import indicated by significant more accumulation of HSPD1 precursor, as well as stabilized PINK1 in the absence of DELE1 pathway compared to wild type. We think this may help to reconcile with the finding that loss of HRI promotes PINK1 stabilization and mitophagy in Hela cells following OA treatment.

Mitochondrial protein import is central to maintaining mitochondrial homeostasis [57]. Understanding how this process is regulated could pave the way for new strategies to modulate mitochondrial function and health. Our studies highlight a novel connection between the ISR, mitochondrial protein import, and mitophagy (Figure 5e). By mildly inhibiting protein synthesis with low doses of cycloheximide or Torin1, an mTOR inhibitor, in ISR-deficient cells, we were able to restore mitochondrial import and suppress mitophagy. Pharmacological inhibition of the ISR with ISRIB or activation of ISR in the absence of DELE1-mediated signaling either enhances or reduces mitophagy, respectively. Additionally, recent research indicates that ATAD1 supports mitochondrial import by extracting clogged precursor proteins resulting from import defects [43]. Future studies should investigate whether ATAD1 works synergistically with the ISR to protect mitochondrial import machinery and suppress mitophagy.

Our findings propose a model in which stressed cells activate the ISR as a "first responder" to sustain mitochondrial import, prioritizing the transport of essential proteins necessary for mitochondrial repair rather than immediately triggering mitophagy. While defects in mitochondrial protein import play a critical role in sensing and responding to mitochondrial stress, they also pose a challenge, as homeostatic proteins must be efficiently imported into mitochondria to perform their repair functions after being upregulated in response to stress. In *C. elegans*, for example, the accumulation of misfolded proteins in mitochondria activates the mitochondrial unfolded protein response (UPR^mt^) by reducing the import efficiency of ATFS1 [63]. When ATFS1 fails to enter the mitochondria, it is relocated to the nucleus, where it acts as a transcription factor to upregulate mitochondrial chaperones and proteases, both of which need to be imported into mitochondria for proper repair. Interestingly, the UPR^mt^ can help preserve mitochondrial protein import by upregulating the mitochondrial import machinery [64]. In mammals, our studies suggest that ISR activation, through inhibition of protein synthesis, can support mitochondrial import under stress. Mitochondrial misfolding stress induced by GTPP (an Hsp90 inhibitor) triggers a mammalian UPR^mt^ by integrating mitochondrial import defects with oxidative stress, leading to the upregulation of mitochondrial homeostasis genes encoding mitochondrial chaperones and proteases [65, 66]. Additionally, GTPP robustly activates the DELE1 axis of the ISR [65, 66]. Although the ISR is not strictly required for UPR^mt^ induction, it is possible that ISR activation enhances mitochondrial import of upregulated mitochondrial chaperons, thereby helping to maintain mitochondrial homeostasis during stress.

## Materials and Methods

### Cell culture and Cell lines

HEK293T cells (ATCC, CRL-3216) and HeLa cells (ATCC, CCL-2) were cultured in Dulbecco’s modified Eagle’s medium (DMEM) (Gibco, 11965-092) with 10% fetal bovine serum (Seradigm, 1300-500), penicillin–streptomycin (Life Technologies, 15140122) and L-glutamine (Life Technologies, 25030081). Cells were cultured at 37°C and 5% CO_2_ in a humidified incubator.

The CRISPRi HEK293T cell line (cXG284) was generated previously. The CRISPRi HeLa cell line was a gift from J. Weissman Lab. The *mtKeima* reporter cell line was generated through lentiviral infection of cXG284 with pMY004 (*CAG:mtKeima*) and FACS-based selection. The *miRFP-PRKN* overexpression cell lines were generated through lentiviral infection of the mtKeima cell line with pXG646 and FACS-based selection. The polyclonal cell line generated after sorting is *PRKN* OE^LL^ cell line. We did monoclonal selection to establish a *PRKN* OE^HL^ expression cell line. Inducible *MTS^COXVIII^-YFP* cell line was generated through lentiviral infection of pXG447 and FACS-based selection after induction by 500 ng/mL Doxycycline overnight.

*ATF4 (KO)* cell lines were generated by CRISPR-Cas9-mediated gene editing by using two sgRNAs targeting the first and second exons of *ATF4*. The protospacer sequences are: 5′-TCACCCTTGCTGTTGTTGGA-3′ and 5′-GCAGAGGATGCCTTCTC-3′ (Synthego, CA). *eIF2α^S49/52/A^* cell lines were generated by CRISPR-Cas9-mediated gene editing through homologous recombination. The HEK293T cells were infected with a sgRNA with protospacer sequence 5′-TTCTTAGTGAATTATCCAGA-3’, a donor ssDNA with a sequence of 5′- GTGAATGTCAGATCCATTGCTGAAATGGGGGCTTATGTCAGCTTGCTGGAATACAACAACATTGAAGGCATGATTCTTCTTGCAGAATTAGCACGAAGACGTATCCGTTCTATCAACAAACTCATCCGAATTGGCAGGAATGAGTGTGTGGTTGTCATTAGGGTGGACAAAGAAAAAGGTAAGTGAGAAAAAT-3’ and Cas9 protein via SNP delivery. *PINK1(KO)* cell lines were generated by CRISPR–Cas9-mediated gene editing by using two sgRNAs targeting the first exon of *PINK1*. The protospacer sequences are: 5′- TCTCCGCTTCTTCCGCCAGT-3′ and 5′-CCTCATCGAGGAAAAACAGG-3′ (cloned into pX459 from Addgene). CRISPRi knockdown cell lines were generated by lentiviral transduction with plasmids containing individual sgRNAs and selected by puromycin at 2 μg/mL. Gene KD or KO was validated through WB or RT-qPCR (Supplementary Figure 7).

Cell lines were tested for mycoplasma contamination routinely with primers oXGmycop-F: 5’TGCACCATCTGTCACTCTGTTAACCTC3’ and oXGmycop-R: 5’GGGAGCAAACAGGATTAGATACCCT3’.

## Constructs

For constructing *mtKeima* reporter (pMY004), *mtKeima* was PCR amplified from pCHAC-mt-mKeima (Addgene, plasmid # 72342) [28] and cloned into a lentiviral vector pdbr3_CAG. For *PRKN* overexpression (pXG646), *PRKN* was PCR amplified from CFP-Parkin (Addgene, plasmid # 47560) [27] and cloned into a lentiviral vector with hPGK promoter and *miRFP*. For inducible *MTS-YFP* (pXG447), mitochondrial targeting sequence of *COXVIII* and *YFP* were cloned from *mito-SRAI* (#RDB18223) [67] into a lentiviral plasmid containing a TetON inducible promoter and trans-activator rtTA. For *PINK1* knock out, sgRNA protospacer sequences were cloned into pSpCas9(BB)-2A- Puro (PX459) V2.0 (Addgene, plasmid # 62988) following the protocol previously published [68]. For gene knockdown via CRISPRi, top and bottom oligonucleotides (IDT) were annealed and ligated to an optimized lentiviral sgRNA expression vector [69].

### Drug treatment

HEK293T cells and its derived cell lines were seeded at 25% confluency 24 hr before drug treatment (i.e., 15,000 cells per well of a 96-well plate). An equal volume of DMEM with twice the final drug concentration was added. For mitochondrial stress, cells were incubated with the following mitochondrial toxins for 24 hr unless otherwise stated: 1.25 ng/ml oligomycin (Sigma-Aldrich, 75351), 100 nM antimycin (Sigma-Aldrich, A8674), 100 nM, rotenone (Sigma-Aldrich, R8875), 10 μM carbonyl cyanide 3- chlorophenylhydrazone (CCCP) (Sigma-Aldrich, C2759), 1.25 ng/ml oligomycin combined with 100 nM antimycin (OA treatment) and 1mM DFP (Sigma-Aldrich, 379409). Bafilomycin A1 (Cayman,11038) was used at a final concentration of 100 nM to suppress autophagy. To attenuate protein synthesis, cycloheximide (Sigma-Aldrich, C7698) was used at a series of concentrations: 50 ng/mL, 100 ng/mL, 200 ng/mL, 400ng/mL, 800ng/mL, 1 μg/mL, 2 μg/mL, 4 μg/mL, 8 μg/mL,10 μg/mL and 20 μg/mL.

mTOR inhibitor Torin1 (Cayman, 10997) was used at concentrations of 50, 100, 200, 250, 500 and 1000 nM. To trigger the GCN2 branch of ISR, halofuginone (Sigma-Aldrich, 50-576-300001) was used at the following concentrations: 2.5, 5, 10, 20, 40, and 80 nM. To activate the PERK branch of ISR, tharpsigargin (Sigma-Aldrich, T9033) was used at the following concentrations: 2.5, 5, 10, 20, 40, and 100 nM.

### Mitophagy measurement by flow cytometry

After 24 hr of drug treatment, cells were digested with trypsin for 3 min at 37°C and then trypsin was neutralized with four to five volumes of DMEM medium. Flow cytometry was performed on an Attune CytPix flow cytometer with an auto sampler (Invitrogen). Events were pre-selected for living and single cell populations. Excitation at 405 nm (pH 7) with 610/20 nm emission filters and excitation at 561 nm (pH 4) with 620/15 nm emission filters were selected to measure mtKeima fluorescence. The flow cytometry data was analyzed using the software FlowJo10.10.0 (FlowJo, LLC). The bar graphs were generated using GraphPad Prism version 10 and Adobe Illustrator.

### Immunoblotting

Cell samples were lysed with RIPA lysate (Thermo Fisher Scientific, Catalog No. 89900) containing protease inhibitor cocktail (Millopore Sigma, Catalog No. 04693159001) for 30 minutes on ice, and vortexed for 15 seconds every 10 minutes. The homogenate was centrifuged at 20,000 × g for 20 min at 4°C, then the supernatant was collected. A BCA kit (Thermo Fisher Scientific, Catalog No. 23227) and a Spark multimode microplate reader (Tecan) was used to measure protein concentration. 20 μg proteins from each sample were separated by 4–12% gradient Bis-Tris Gels (Thermo Fisher Scientific) and transferred to 0.2 μm nitrocellulose membranes (Thermo Fisher Scientific, Catalog No. 77012). The membrane was blocked with 5% nonfat milk diluted in TBS (20 mM Tris-HCl, 150 mM NaCl, pH 7.5) for 1 hr with gentle shaking at room temperature (RT), then incubated with the primary antibodies overnight (about 15 hr) at 4°C. After three washes with TBS-T (TBS containing 0.1% Tween-20), the membrane was incubated with infrared fluorescent dye labeled secondary antibodies (LI-COR) for 2 hr at RT with gentle shaking, followed by washing with TBS-T three times for 10 min each at RT. Images were visualized by an Odyssey imager (LI-COR) and processed using ImageStudio Lite v5.2 (LI-COR). Antibodies used in this study include: anti-ATF4 (ProteinTech, 28657-1-AP, rabbit, 1:1000), anti-β-actin (ProteinTech, 66009-1-Ig, mouse, 1:5000), anti-β-actin (ProteinTech, 81115-1-RR, rabbit, 1:5000), anti-COXIV (Invitrogen, MA5-17279, mouse, 1:2000), anti-eIF2α (ProteinTech, 11170-1-AP, Rabbit, 1:1000), anti-GFP (Roche, 11814460001, mouse, 1:1,000), anti-HSPD1 (Invitrogen, MA3-012, mouse, 1:2000), anti-LC3 (ProteinTech, 14600-1-AP, Rabbit, 1:500), anti-phospho-eIF2α(ser51) (Cell Signaling Technology, 3398S, Rabbit, 1:1000), anti-PINK1 (ProteinTech, 23274-1-AP, rabbit, 1:500), anti-puromycin (Millipore Sigma, MABE343, mouse, 1:20000), anti-VDAC (Cell Signaling Technology, 4661S, Rabbit, 1:1000)

### Puromycin labeling assay

Cells were seeded at 25% confluency in 10-cm dishes (Thermo-Scientific, #130182) 24 h before drug treatment. To depolarize the mitochondrial membrane potential, cells were treated with 10 μM CCCP (Sigma-Aldrich, C2759) for 2.5 hr as a positive control. For mitochondrial stress, cells were incubated with 2.5 ng/ml oligomycin for 2.5 hr (Sigma-Aldrich, 75351). Before harvesting, cells were treated with 10 μg/ml puromycin (Sigma-Aldrich, P4512) for 0.5 h to label newly synthesized proteins. Cells were harvested and washed with D-PBS (Gibco, 2031651). 10% of the cells were split to a separate tube to measure total protein synthesis and ATF4 induction. The remaining cells were used to isolate mitochondria using a mitochondria isolation kit (Thermo-Scientific, YK380589). Cells were separated into two proportions (cytosol and mitochondria) during this process, and mitochondrial protein import efficiency was evaluated through immunoblotting of puromycin using a anti-puromycin antibody (Millipore Sigma, MABE343, mouse, 1:20000).

### Mitochondrial membrane potential measurement

Cells were seeded at 25% confluency in 24-well plates (Thermo-Scientific, 930186) 24 hr before drug treatment. To depolarize mitochondrial membrane, cells were treated with 10 μM CCCP (Sigma-Aldrich, C2759) for 3 hr as a positive control. For mitochondrial stress, cells were incubated with 1.25 ng/ml oligomycin (Sigma-Aldrich, 75351) 3 hr and 24 hr. To measure mitochondrial membrane potential, cells were stained with 100 nM tetramethylrhodamine (TMRE) (Invitrogen, T669) in DMEM media for 20 minutes at 37°C. After TMRE staining, cells were harvested by trypsinization and neutralized with DMEM (Gibco, 2910058) media, then washed twice with D-PBS (with 0.5 % FBS). Cells were flowed using Attune CytPix flow cytometer. TMRE signal was detected through a 561 nm laser and 574 nm emission filter. Recorded data was analyzed using FlowJo software. Cells were gated by forward and side scatter for viable, single cells. BFP signals indicate the cells containing sgRNA for CRISPRi.

### Live cell imaging

Cells were seeded at 25% confluence on 8-chamber glass slides (Cellvis, C8-1.5H-N). For induction of MTS-YFP during mitochondrial stress, cells were incubated with 500 ng/mL doxycycline (Sigma-Aldrich, 631311) and 1.25 ng/ml oligomycin (Sigma-Aldrich, 75351) overnight. Cells were stained with 100 nM MitoTracker Red (Invitrogen, 2549283) for 0.5 hr before imaging. Imaging was performed with a Zeiss 780 confocal microscope (Carl Zeiss) equipped with a 63× oil immersion objective. Images were obtained using the ZEN software (Carl Zeiss). Within every experiment, laser percentage and exposure time for each channel were set the same across all samples in comparison. All images were analyzed using Fiji.

### RT-qPCR

Total RNA was extracted using the Quick-RNA Miniprep Kit (Zymo Research, R1055), and first-strand cDNA was synthesized with Maxima H Minus First Strand cDNA Synthesis Kit (Thermo Scientific™, K1652). qPCR was performed with Applied Biosystems™ PowerUp™ SYBR™ Green Master Mix for qPCR (Applied Biosystems™, A25742). Fold changes in expression were calculated using the ΔΔCt method. qPCR primers used are as following: Actin_F: 5’-GTCATCACCATTGGCAATGAG-3’ Actin_R: 5’-CGTCATACTCCTGCTTGCTG-3’ ASNS_F: 5’-ATCACTGTCGGGATGTACCC-3’ ASNS_R: 5’-TGATAAAAGGCAGCCAATCC-3’ DDIT3_F: 5’-AGCCAAAATCAGAGCTGGAA-3’ DDIT3_R: 5’-TGGATCAGTCTGGAAAAGCA-3’ HRI_F: 5’-ACACCAACACATACGTCCAG-3’ HRI_R: 5’-GCTCCATTTCTGTTCCAAACG-3’ DELE1_F: 5’-AGGCTGTGACTTCCATTCAG-3’ DELE1_F: 5’-TCGCCACTCTTCATGTTCTC-3’

## Acknowledgements

We thank Dr. Ruilin Tian, Dr. Emmy Li and Dr. Bernard Cook for critical reading of this manuscript. We thank Dr. Evan Jellison and Li Zhu from the UConn Health Flow Cytometry Facility for their technical support on cell sorting. We thank Dr. Yi Wu and Susan Staurovsky from the CCAM Microscopy Facility for their technical support on imaging. This work was funded by the startup from UConn Health (XG), the Research Excellence Program from University of Connecticut (XG) and the National institutes of Health (R35GM155240 to XG).

## Author contributions

MY, ZM, and XG conceived, designed, performed, and interpreted the experiments. KW and WL performed experiments. XG supervised the project. MY, ZM, and XG wrote the manuscript.

---

**Supplementary Figure 1.**
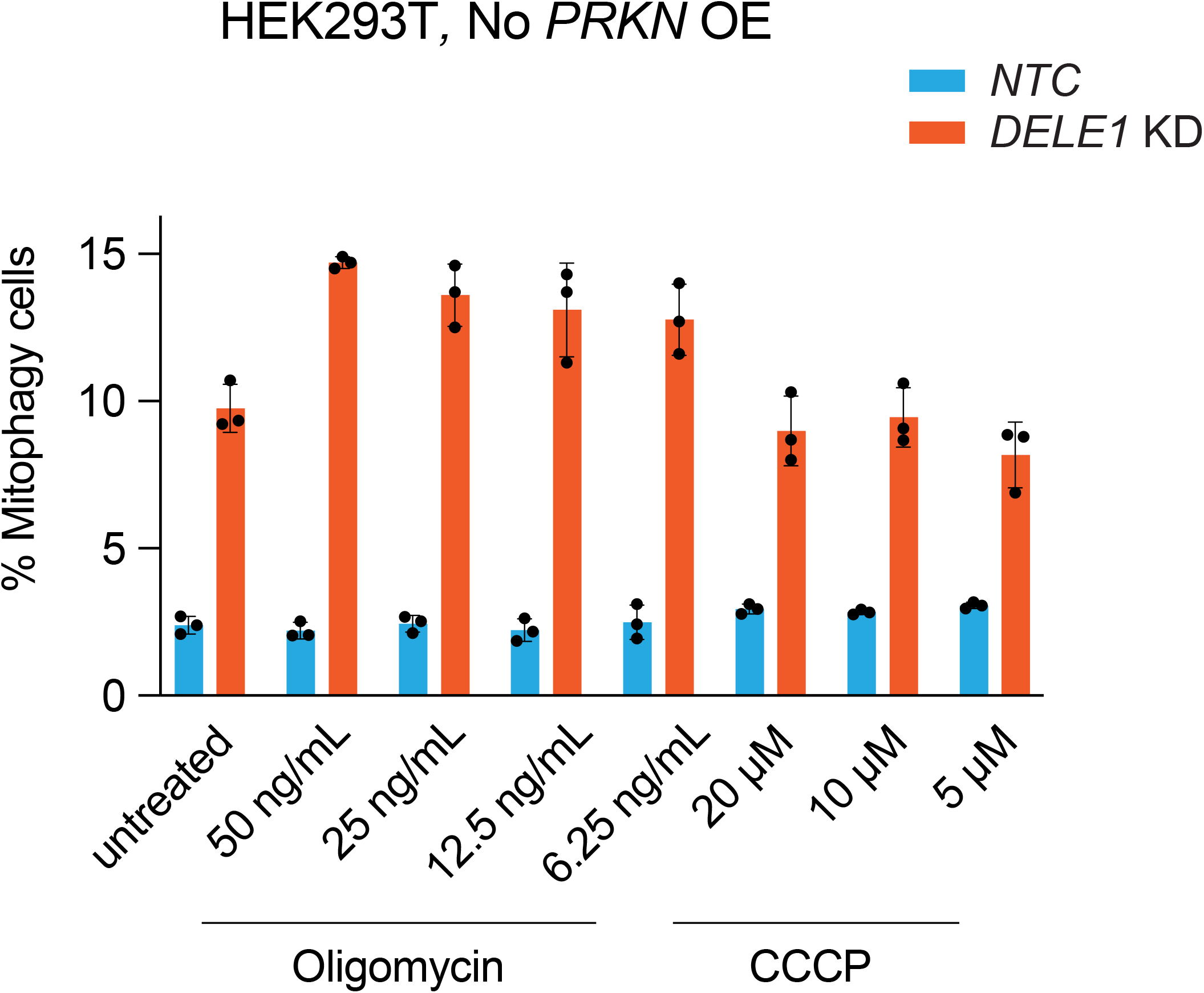
Mitophagy level in HEK293T cells without overexpression of PRKN. Mitophagy was measured in *NTC* and *DELE1* KD cells without *PRKN* overexpression were treated with oligomycin at concentrations from 6.25 to 50 ng/mL, and CCCP at concentrations from 5 to 20 μM.

**Supplementary Figure 2.**
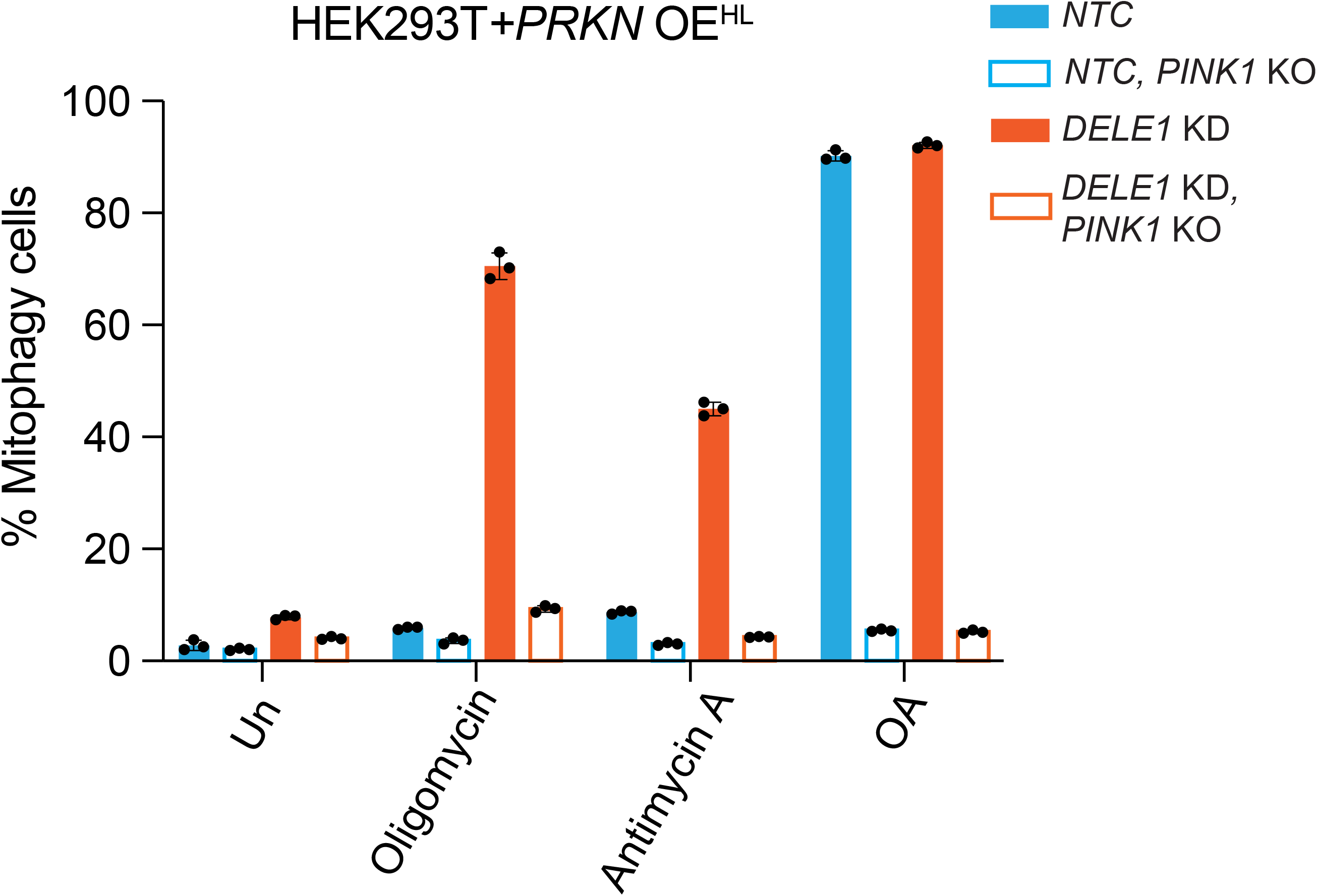
Regulation of mitophagy under oligomycin plus antimycin A (OA) in HEK293T cells. HEK293T *PRKN* OE^HL^ wild type (*WT*) or *PINK1 (KO)* cells with or without *DELE1* KD were treated with 1.25 ng/ml oligomycin, 100 nM antimycin, or a combination of oligomycin and antimycin A (OA) for 24 hr, followed by measurement of mitophagy using flow cytometry. (mean ± s.d., n = 3 culture wells)

**Supplementary Figure 3.**
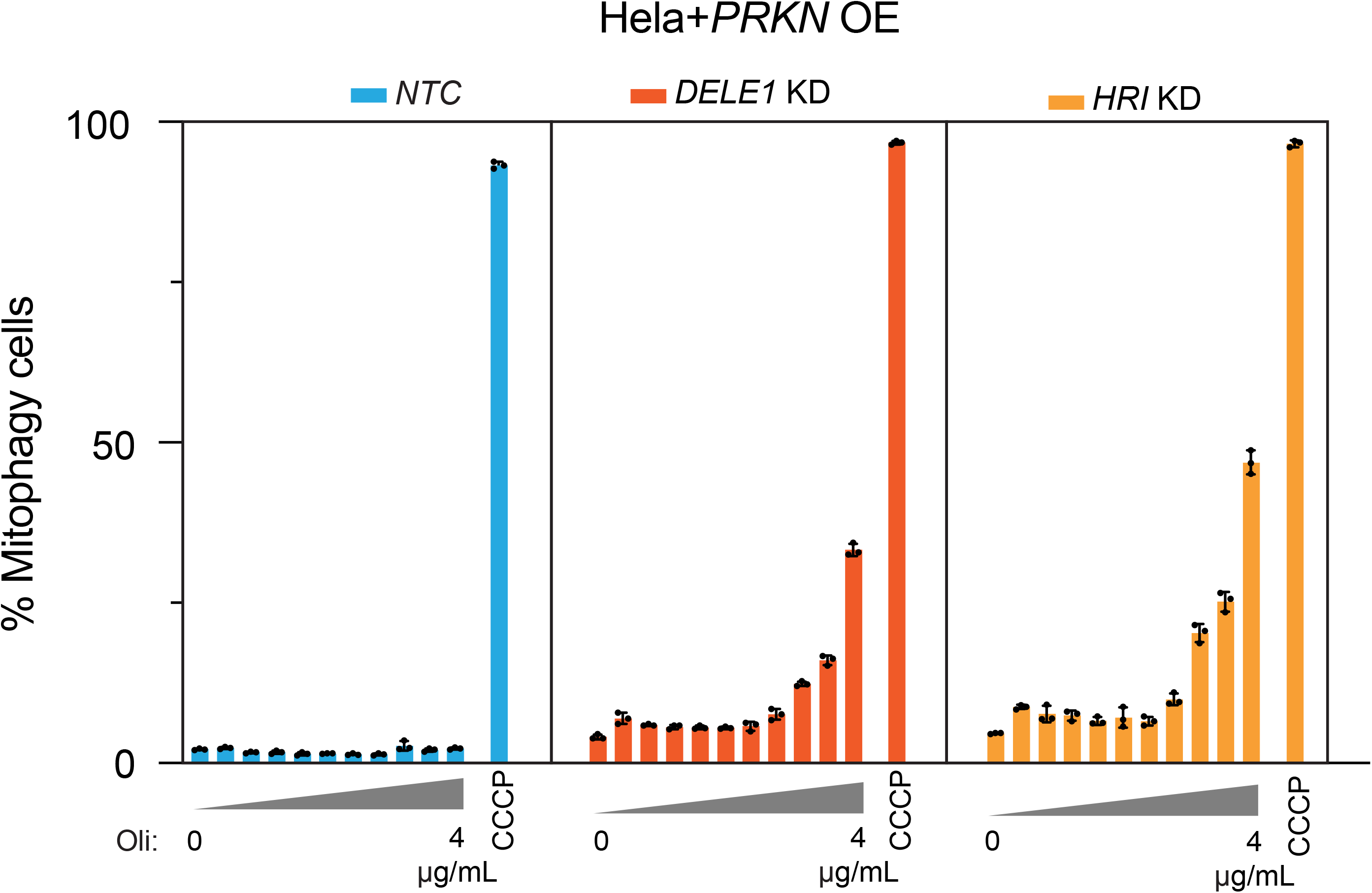
DELE1 pathway negatively regulates mitophagy in Hela cells. Hela cell line with *PRKN* OE were infected with *NTC* sgRNA, *DELE1* sgRNA or *HRI* sgRNA. These cells were treated with oligomycin (Oli) at following concentrations: 1.25 ng/mL, 2.5 ng/mL, 5 ng/mL, 10 ng/mL, 50 ng/mL, 100 ng/mL, 500 ng/mL, 1 μg/mL, 2 μg/mL, and 4 μg/mL, as well as CCCP at 10 μM for 24 hr, followed by flow cytometry measurement of mitophagy.

**Supplementary Figure 4.**
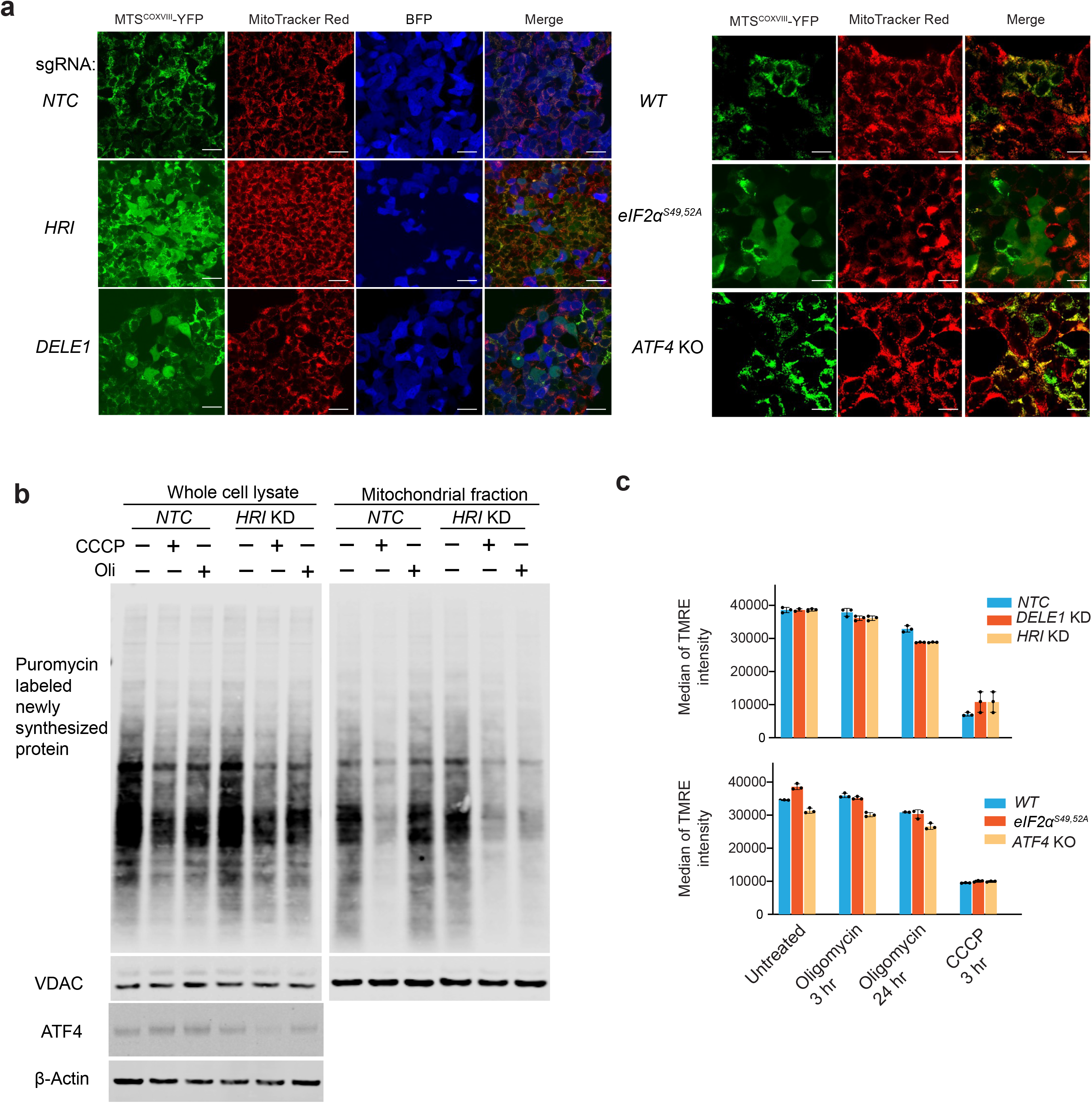
DELE1-ISR preserves mitochondrial protein import. **(a)** Inducible *MTS^COXVIII^-YFP* was integrated into cells with *NTC* sgRNA, *HRI* sgRNA, *DELE1* sgRNA, wild type (*WT*), *eIF2α^S49/52/A^* and *ATF4* KO. These cells were treated with 500 ng/mL doxycycline and 1.25 ng/mL oligomycin for overnight before imaging. MitoTracker Red stains mitochondria. Scale bar: 20 μm. **(b)** Immunoblot of puromycin in isolated mitochondria and whole cell lysate. Cells with *NTC* or *HRI* sgRNAs were treated with 10 μM CCCP and 1.25 ng/mL oligomycin for 3 hr. ATF4 activation indicates mitochondrial stress and *HRI* knockdown. β-Actin (whole cell) and VDAC (mitochondrial fraction) serve as the loading control. **(c)** HEK293T cells with *NTC* sgRNA, *HRI* sgRNA, *DELE1* sgRNA, wild type (*WT*), *eIF2α^S49/52/A^* and *ATF4* KO were treated with 10 μM CCCP for 3 hr, as a positive control for mitochondrial depolarization, and 1.25 ng/ml oligomycin for 3 or 24 hr, followed by 100 nM TMRE staining and flow cytometry analysis. (mean ± s.d., n = 3 culture wells)

**Supplementary Figure 5.**
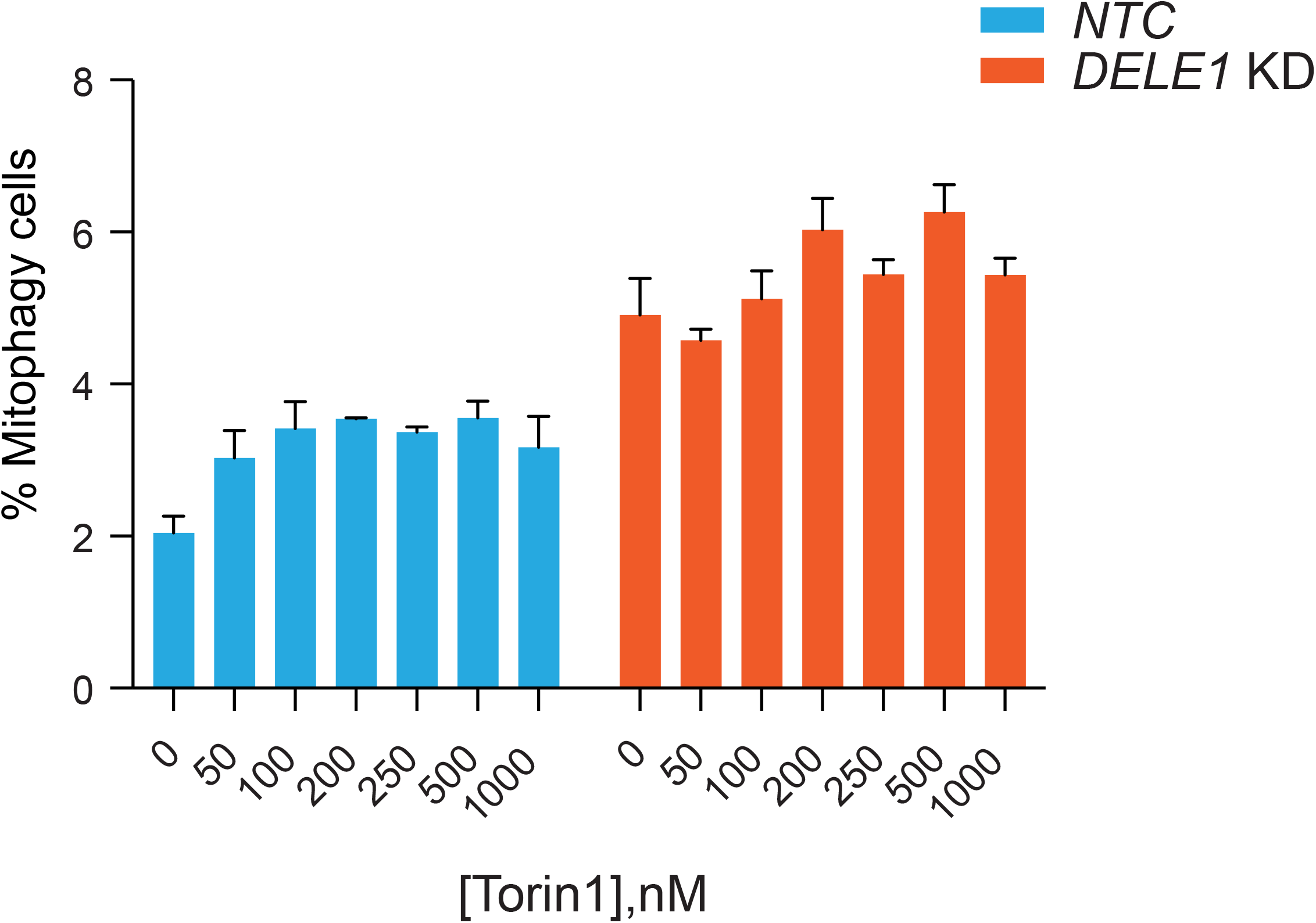
Torin1 slightly increases mitophagy. *NTC* and *DELE1* KD cells were treated with torin1 at concentrations from 50 nM to 1000 nM, followed by flow cytometry to measure mitophagy. (mean ± s.d., n = 3 culture wells)

**Supplementary Figure 6.**
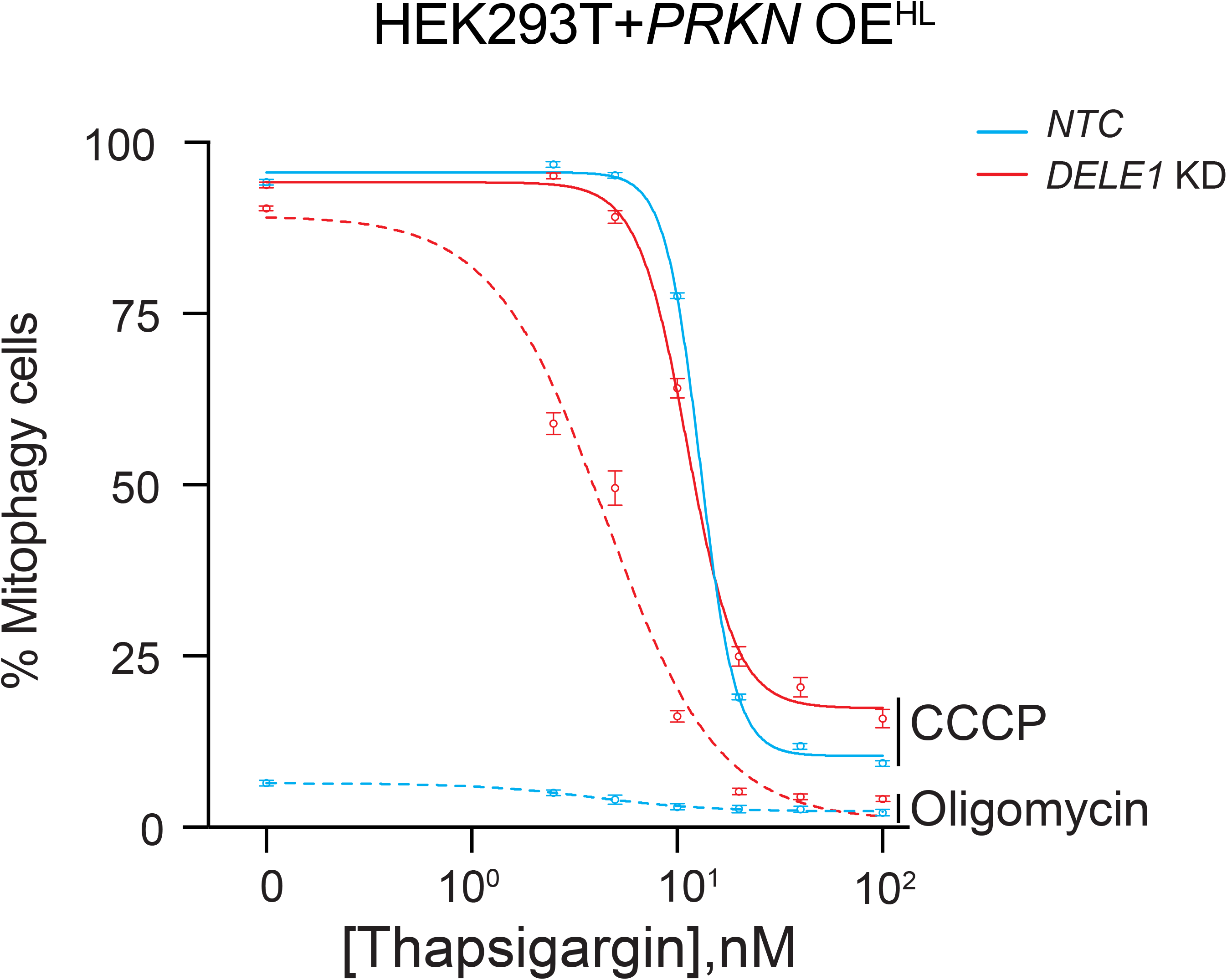
Activation of PERK-ISR by tharpsigargin suppress mitophagy in DELE1-ISR-deficient cells. *NTC* and *DELE1* KD cells with *PRKN* OE^HL^ are treated with 10 μM CCCP or 1.25 ng/mL oligomycin in the presence of tharpsigargin at 7 different concentrations (0, 2.5 nM, 5 nM, 10 nM, 20 nM, 40 nM and 100 nM) for 24 hr followed by flow cytometry to measure mitophagy. Tharpsigargin concentrations were converted to their base-10 logarithmic values. A nonlinear regression analysis using a log(inhibitor) vs. response model with a variable slope (four parameters) was performed to generate the plot. (mean ± s.d., n = 3 culture wells)

**Supplementary Figure 7.**
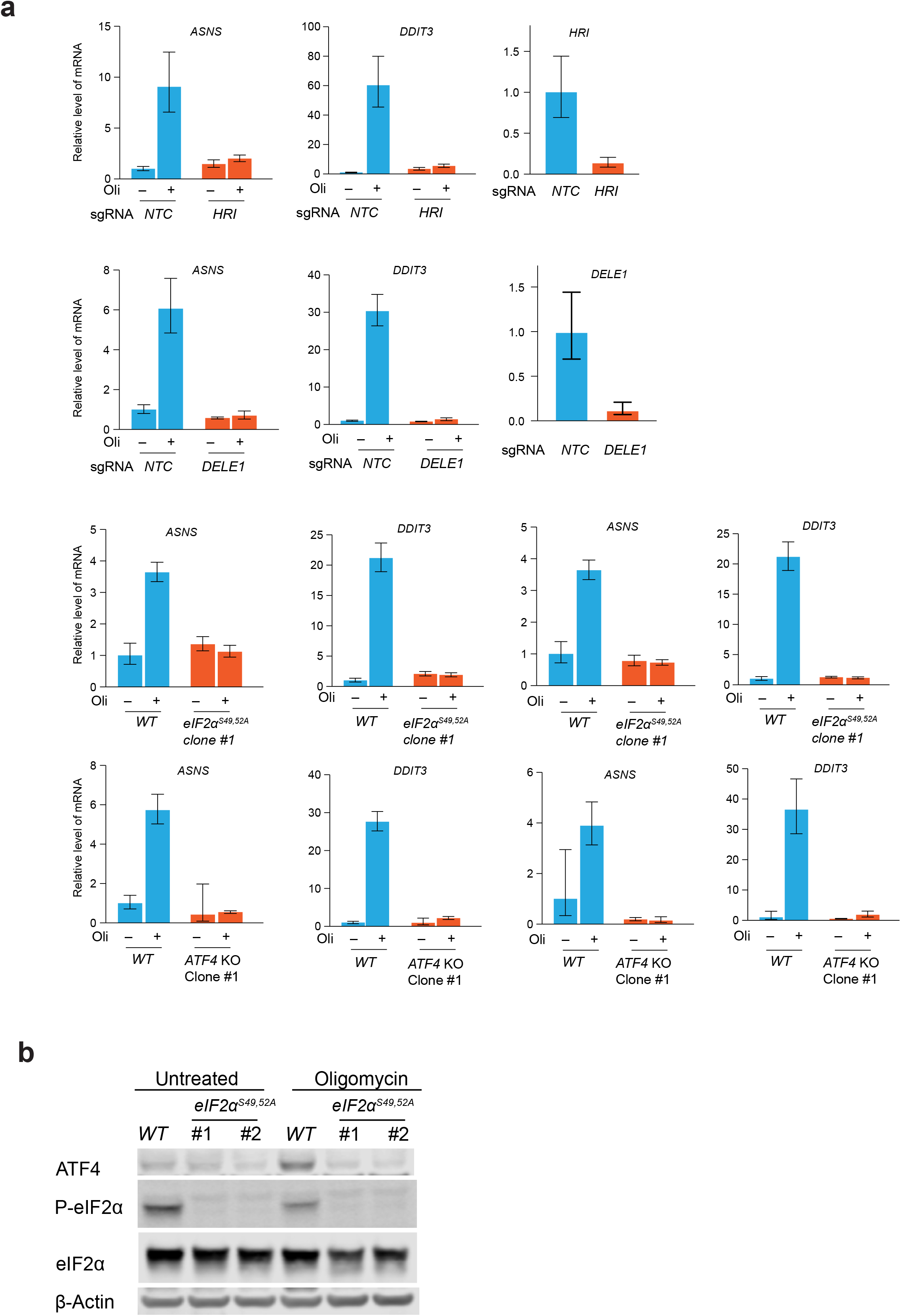
Validation of loss of the ISR activation in different cell lines. **(a)** *NTC, HRI* KD and *DELE1* KD, wild type (*WT*), *eIF2α^S49/52/A^* and *ATF4* KO cells were left untreated or treated with 1.25 ng/mL oligomycin for 6 hr before harvesting for RNA extraction, cDNA synthesis and qPCR. ISR target genes *ASNS* and *DDIT3* were used to evaluate the activation of the ISR. For *DELE1* and *HRI* knockdown, qPCR of *DELE1* and *HRI* were also performed. **(b)** Immunoblots of ATF4, phosphorylated eIF2α, total eIF2α and β-actin. *WT* and two clonal *eIF2α^S49/52/A^* cells were treated with 1.25 ng/mL oligomycin for 24 hr. β-Actin serves as the loading control.

## Reference

1. Suomalainen, A. and J. Nunnari, Mitochondria at the crossroads of health and disease. Cell, 2024. 187(11): p. 2601–2627.

2. Picard, M. and O.S. Shirihai, Mitochondrial signal transduction. Cell Metab, 2022. 34(11): p. 1620–1653.

3. Bao, X.R., et al., Mitochondrial dysfunction remodels one-carbon metabolism in human cells. Elife, 2016. 5.

4. Quiros, P.M., et al., Multi-omics analysis identifies ATF4 as a key regulator of the mitochondrial stress response in mammals. J Cell Biol, 2017. 216(7): p. 2027–2045.

5. Costa-Mattioli, M. and P. Walter, The integrated stress response: From mechanism to disease. Science, 2020. 368(6489).

6. Adomavicius, T., et al., The structural basis of translational control by eIF2 phosphorylation. Nat Commun, 2019. 10(1): p. 2136.

7. Kashiwagi, K., et al., Structural basis for eIF2B inhibition in integrated stress response. Science, 2019. 364(6439): p. 495–499.

8. Kenner, L.R., et al., eIF2B-catalyzed nucleotide exchange and phosphoregulation by the integrated stress response. Science, 2019. 364(6439): p. 491–495.

9. Fessler, E., et al., A pathway coordinated by DELE1 relays mitochondrial stress to the cytosol. Nature, 2020. 579(7799): p. 433–437.

10. Guo, X., et al., Mitochondrial stress is relayed to the cytosol by an OMA1-DELE1-HRI pathway. Nature, 2020. 579(7799): p. 427–432.

11. Sekine, Y., et al., A mitochondrial iron-responsive pathway regulated by DELE1. Mol Cell, 2023. 83(12): p. 2059–2076 e6.

12. Yang, J., et al., DELE1 oligomerization promotes integrated stress response activation. Nat Struct Mol Biol, 2023. 30(9): p. 1295–1302.

13. Palikaras, K., E. Lionaki, and N. Tavernarakis, Mechanisms of mitophagy in cellular homeostasis, physiology and pathology. Nat Cell Biol, 2018. 20(9): p. 1013–1022.

14. Pickles, S., P. Vigie, and R.J. Youle, Mitophagy and Quality Control Mechanisms in Mitochondrial Maintenance. Curr Biol, 2018. 28(4): p. R170–R185.

15. Pickrell, A.M. and R.J. Youle, The roles of PINK1, parkin, and mitochondrial fidelity in Parkinson’s disease. Neuron, 2015. 85(2): p. 257–73.

16. Kitada, T., et al., Mutations in the parkin gene cause autosomal recessive juvenile parkinsonism. Nature, 1998. 392(6676): p. 605–8.

17. Valente, E.M., et al., Hereditary early-onset Parkinson’s disease caused by mutations in PINK1. Science, 2004. 304(5674): p. 1158–60.

18. Narendra, D.P. and R.J. Youle, The role of PINK1-Parkin in mitochondrial quality control. Nat Cell Biol, 2024.

19. Deas, E., et al., PINK1 cleavage at position A103 by the mitochondrial protease PARL. Hum Mol Genet, 2011. 20(5): p. 867–79.

20. Meissner, C., et al., The mitochondrial intramembrane protease PARL cleaves human Pink1 to regulate Pink1 trafficking. J Neurochem, 2011. 117(5): p. 856–67.

21. Jin, S.M., et al., Mitochondrial membrane potential regulates PINK1 import and proteolytic destabilization by PARL. J Cell Biol, 2010. 191(5): p. 933–42.

22. Yamano, K. and R.J. Youle, PINK1 is degraded through the N-end rule pathway. Autophagy, 2013. 9(11): p. 1758–69.

23. Narendra, D.P., et al., PINK1 is selectively stabilized on impaired mitochondria to activate Parkin. PLoS Biol, 2010. 8(1): p. e1000298.

24. Vives-Bauza, C., et al., PINK1-dependent recruitment of Parkin to mitochondria in mitophagy. Proc Natl Acad Sci U S A, 2010. 107(1): p. 378–83.

25. Chan, N.C., et al., Broad activation of the ubiquitin-proteasome system by Parkin is critical for mitophagy. Hum Mol Genet, 2011. 20(9): p. 1726–37.

26. Sarraf, S.A., et al., Landscape of the PARKIN-dependent ubiquitylome in response to mitochondrial depolarization. Nature, 2013. 496(7445): p. 372–6.

27. Narendra, D., et al., Parkin is recruited selectively to impaired mitochondria and promotes their autophagy. J Cell Biol, 2008. 183(5): p. 795–803.

28. Lazarou, M., et al., The ubiquitin kinase PINK1 recruits autophagy receptors to induce mitophagy. Nature, 2015. 524(7565): p. 309–314.

29. Heo, J.M., et al., The PINK1-PARKIN Mitochondrial Ubiquitylation Pathway Drives a Program of OPTN/NDP52 Recruitment and TBK1 Activation to Promote Mitophagy. Mol Cell, 2015. 60(1): p. 7–20.

30. Wong, Y.C. and E.L. Holzbaur, Optineurin is an autophagy receptor for damaged mitochondria in parkin-mediated mitophagy that is disrupted by an ALS-linked mutation. Proc Natl Acad Sci U S A, 2014. 111(42): p. E4439–48.

31. Michaelis, J.B., et al., Protein import motor complex reacts to mitochondrial misfolding by reducing protein import and activating mitophagy. Nat Commun, 2022. 13(1): p. 5164.

32. Sidrauski, C., et al., Pharmacological brake-release of mRNA translation enhances cognitive memory. Elife, 2013. 2: p. e00498.

33. Sidrauski, C., et al., Pharmacological dimerization and activation of the exchange factor eIF2B antagonizes the integrated stress response. Elife, 2015. 4: p. e07314.

34. Katayama, H., et al., A sensitive and quantitative technique for detecting autophagic events based on lysosomal delivery. Chem Biol, 2011. 18(8): p. 1042–52.

35. Qi, L.S., et al., Repurposing CRISPR as an RNA-guided platform for sequence-specific control of gene expression. Cell, 2013. 152(5): p. 1173–83.

36. Loomis, W.F. and F. Lipmann, Reversible inhibition of the coupling between phosphorylation and oxidation. J Biol Chem, 1948. 173(2): p. 807.

37. Kabeya, Y., et al., LC3, a mammalian homologue of yeast Apg8p, is localized in autophagosome membranes after processing. Embo j, 2000. 19(21): p. 5720–8.

38. Waters, C.S., et al., A PINK1 input threshold arises from positive feedback in the PINK1/Parkin mitophagy decision circuit. Cell Rep, 2023. 42(10): p. 113260.

39. Yang, H., et al., Dynamic Modeling of Mitochondrial Membrane Potential Upon Exposure to Mitochondrial Inhibitors. Front Pharmacol, 2021. 12: p. 679407.

40. Jin, S.M. and R.J. Youle, The accumulation of misfolded proteins in the mitochondrial matrix is sensed by PINK1 to induce PARK2/Parkin-mediated mitophagy of polarized mitochondria. Autophagy, 2013. 9(11): p. 1750–7.

41. Schmidt, E.K., et al., SUnSET, a nonradioactive method to monitor protein synthesis. Nat Methods, 2009. 6(4): p. 275–7.

42. Martensson, C.U., et al., Mitochondrial protein translocation-associated degradation. Nature, 2019. 569(7758): p. 679–683.

43. Kim, J., et al., ATAD1 prevents clogging of TOM and damage caused by un-imported mitochondrial proteins. Cell Rep, 2024. 43(8): p. 114473.

44. Thoreen, C.C., et al., An ATP-competitive mammalian target of rapamycin inhibitor reveals rapamycin-resistant functions of mTORC1. J Biol Chem, 2009. 284(12): p. 8023–32.

45. Morita, M., et al., mTORC1 Controls Mitochondrial Activity and Biogenesis through 4E-BP-Dependent Translational Regulation. Cell Metabolism, 2013. 18(5): p. 698–711.

46. Park, Y., et al., mTORC1 Balances Cellular Amino Acid Supply with Demand for Protein Synthesis through Post-transcriptional Control of ATF4. Cell Rep, 2017. 19(6): p. 1083–1090.

47. Perea, V., et al., Pharmacologic activation of a compensatory integrated stress response kinase promotes mitochondrial remodeling in PERK-deficient cells. Cell Chem Biol, 2023. 30(12): p. 1571–1584 e5.

48. Keller, T.L., et al., Halofuginone and other febrifugine derivatives inhibit prolyl-tRNA synthetase. Nat Chem Biol, 2012. 8(3): p. 311–7.

49. Thastrup, O., et al., Thapsigargin, a tumor promoter, discharges intracellular Ca2+stores by specific inhibition of the endoplasmic reticulum Ca2(+)-ATPase. Proc Natl Acad Sci U S A, 1990. 87(7): p. 2466–70.

50. Bertolotti, A., et al., Dynamic interaction of BiP and ER stress transducers in the unfolded-protein response. Nat Cell Biol, 2000. 2(6): p. 326–32.

51. Ahola, S., et al., OMA1-mediated integrated stress response protects against ferroptosis in mitochondrial cardiomyopathy. Cell Metab, 2022. 34(11): p. 1875–1891 e7.

52. Huynh, H., et al., DELE1 is protective for mitochondrial cardiomyopathy. J Mol Cell Cardiol, 2023. 175: p. 44–48.

53. Zhu, S., et al., Mitochondrial Stress Induces an HRI-eIF2alpha Pathway Protective for Cardiomyopathy. Circulation, 2022. 146(13): p. 1028–1031.

54. Lin, H.P., et al., DELE1 maintains muscle proteostasis to promote growth and survival in mitochondrial myopathy. EMBO J, 2024.

55. Chakrabarty, Y., et al., The HRI branch of the integrated stress response selectively triggers mitophagy. Mol Cell, 2024. 84(6): p. 1090–1100 e6.

56. Singh, P.K., et al., Kinome screening identifies integrated stress response kinase EIF2AK1 / HRI as a negative regulator of PINK1 mitophagy signaling. bioRxiv, 2024: p. 2023.03.20.533516.

57. Lionaki, E., I. Gkikas, and N. Tavernarakis, Mitochondrial protein import machinery conveys stress signals to the cytosol and beyond. Bioessays, 2023. 45(3): p. e2200160.

58. Haastrup, M.O., et al., The Journey of Mitochondrial Protein Import and the Roadmap to Follow. Int J Mol Sci, 2023. 24(3).

59. Fessler, E., L. Krumwiede, and L.T. Jae, DELE1 tracks perturbed protein import and processing in human mitochondria. Nat Commun, 2022. 13(1): p. 1853.

60. Bi, P.Y., et al., Cytosolic retention of HtrA2 during mitochondrial protein import stress triggers the DELE1-HRI pathway. Commun Biol, 2024. 7(1): p. 391.

61. Shpilka, T., et al., UPR(mt) scales mitochondrial network expansion with protein synthesis via mitochondrial import in Caenorhabditis elegans. Nat Commun, 2021. 12(1): p. 479.

62. Fu, Y., et al., Mitochondrial DNA breaks activate an integrated stress response to reestablish homeostasis. Mol Cell, 2023. 83(20): p. 3740–3753 e9.

63. Nargund, A.M., et al., Mitochondrial import efficiency of ATFS-1 regulates mitochondrial UPR activation. Science, 2012. 337(6094): p. 587–90.

64. Xin, N., et al., The UPRmt preserves mitochondrial import to extend lifespan. J Cell Biol, 2022. 221(7).

65. Sutandy, F.X.R., et al., A cytosolic surveillance mechanism activates the mitochondrial UPR. Nature, 2023. 618(7966): p. 849–854.

66. Munch, C. and J.W. Harper, Mitochondrial unfolded protein response controls matrix pre-RNA processing and translation. Nature, 2016. 534(7609): p. 710–3.

67. Katayama, H., et al., Visualizing and Modulating Mitophagy for Therapeutic Studies of Neurodegeneration. Cell, 2020. 181(5): p. 1176–1187 e16.

68. Ran, F.A., et al., Genome engineering using the CRISPR-Cas9 system. Nat Protoc, 2013. 8(11): p. 2281–2308.

69. Gilbert, L.A., et al., Genome-Scale CRISPR-Mediated Control of Gene Repression and Activation. Cell, 2014. 159(3): p. 647–61.

